# Her9/Hes4 is required for retinal photoreceptor development, maintenance, and survival

**DOI:** 10.1101/833301

**Authors:** Cagney E. Coomer, Stephen G. Wilson, Kayla F. Titialii-Torres, Jessica D. Bills, Laura A. Krueger, Rebecca A. Petersen, Evelyn M. Turnbaugh, Eden L. Janesch, Ann C. Morris

## Abstract

The intrinsic and extrinsic factors that regulate vertebrate photoreceptor specification and differentiation are complex, and our understanding of all the players is far from complete. Her9, the zebrafish ortholog of human HES4, is a basic helix-loop-helix-orange (bHLH-O) transcriptional repressor that regulates neurogenesis in several developmental contexts. We have previously shown that *her9* is upregulated during chronic rod photoreceptor degeneration and regeneration in adult zebrafish, but little is known about the role of *her9* during retinal development. To better understand the function of Her9 in the retina, we generated zebrafish *her9* CRISPR mutants. *Her9* homozygous mutants displayed striking retinal phenotypes, including decreased numbers of rods and red/green cones, whereas blue and UV cones were relatively unaffected. The reduction in rods and red/green cones correlated with defects in photoreceptor subtype lineage specification. The remaining rods and double cones displayed abnormally truncated outer segments, and elevated levels of apoptosis. In addition to the photoreceptor defects, *her9* mutants also possessed a reduced proliferative ciliary marginal zone, and decreased and disorganized Müller glia. Mutation of *her9* was larval lethal, with no mutants surviving past 13 days post fertilization. Our results reveal a previously undescribed role for Her9/Hes4 in photoreceptor differentiation, maintenance, and survival.

## Introduction

The vertebrate retina is a highly conserved tissue of the central nervous system (CNS) that captures and converts light into an electrical signal. The neural retina is composed of three layers; the ganglion cell layer (GCL), inner nuclear layer (INL) and outer nuclear layer (ONL); the light-sensitive rod and cone photoreceptors reside in the ONL. The six classes of retinal neurons and single glial cell type in the retina all differentiate from a single pool of multipotent retinal progenitor cells (RPCs). Across all vertebrates studied to date, specification and differentiation of retinal cell types occurs in a synchronized and largely conserved manner. The ganglion cells are born first and the rods, bipolar cells, and Müller glia differentiate last (Livesey and Cepko, 2001). Cone photoreceptor differentiation generally occurs before the rods. The development of photoreceptor cell identity is influenced by many factors, both intrinsic and extrinsic. Cone and rod photoreceptor specification, cell cycle exit, and differentiation are accompanied by the expression of photoreceptor-specific transcription factors such as *otx2* and *crx*, then rod or cone-specific factors such as *Nr2e3* and *trβ2* (Farre et al., 2019; Il et al., 2010). Photoreceptor differentiation is also regulated extrinsically by Hedgehog (Hh), thyroid hormone (TH), Notch, Wnt, retinoic acid (RA), and fibroblast growth factor (FGF) signaling pathways, to name a few (Doerre and Malicki, 2002). Our knowledge of the precise molecular determinants of photoreceptor subtype differentiation is far from complete. Understanding how photoreceptor cell development is regulated is essential for understanding the pathogenesis of blinding retinal degenerative diseases such as retinitis pigmentosa and macular degeneration. Several approaches have been developed to understand retinal development, one of the most prominent being the generation and characterization of genetic mutants. Due to its conserved retinal structure and function, as well as the extensive genetic homology with humans, zebrafish have become an appealing model for interrogating the role of specific genes in retinal cell type differentiation.

In this study, we examine the role of the transcription factor *Hairy and enhancer of split related 9* (Her9) in retinal development. Her genes belong to a family of basic-Helix-Loop-Helix-Orange (bHLH-O) DNA binding proteins (Muller et al., 1996), and are homologs of the *Hairy* and *Enhancer-of-split* genes in Drosophila and of *Hes/Hey* genes in mammals. The zebrafish genome contains 19 *hairy-related (her)* genes which have been shown to play essential roles in developmental processes such as somitogenesis, neural tube and nervous system development, floor plate development, cell cycle exit, and apoptosis (Liu et al., 2015; Takke et al., 1999). Her proteins often function as downstream effectors of the Notch-Delta signaling pathway and mediate cross-talk between Notch and other pathways (Delidakis et al., 2014). A variety of members in the three *hairy* classes are also regulated by Hh, VEGF, Nodal, FGF, and RA signaling (Holzschuh et al., 2005a; Holzschuh et al., 2005b; Latimer et al., 2005; Morrow et al., 2009; Radosevic et al., 2011).

Zebrafish Her9 shares homology with mouse and human Hes1 (Radosevic et al., 2011), but is more closely related by sequence comparison and synteny analysis to human HES4 (El Yakoubi et al., 2012). As Hes4 is absent from the mouse and rat genomes, zebrafish provide an unique opportunity to explore the function of this transcription factor during retinal development. In zebrafish, *her9* is expressed in ectodermal tissues (Leve et al., 2001) and in inter-pro-neural domains during embryonic development (Bae et al., 2005), where morpholino knockdown suggested it acts downstream of Bmp signaling. Recent studies have also shown that Her9 functions during inner ear development, where it defines the posterolateral non-neurogenic field downstream of *tbx1* expression and RA signaling (Radosevic et al., 2011). Interestingly, *her9* expression was shown to be independent of Notch-Delta signaling in several developmental contexts (Bae et al., 2005; Leve et al., 2001), suggesting that it does not function as a classical Notch effector gene. In the post-embryonic Medaka and Xenopus retina, Her9 regulates proliferation of retinal stem cells in the peripheral ciliary marginal zone (CMZ) (El Yakoubi et al., 2012; Reinhardt et al., 2015). Finally, *her9* expression is upregulated during the specific degeneration and regeneration of rod photoreceptors in the zebrafish retina, suggesting a role for this transcription factor in retinal regeneration (Morris et al., 2011). Although these studies strongly implicate *her9* in regulating continual neurogenesis and injury-induced regeneration in the post-embryonic retina, the function of *her9* during embryonic retinal development has not been thoroughly investigated.

In this study, we generated *her9* mutants using the CRISPR/Cas9 system to determine whether *her9* has a required role in retinal development. Here, we demonstrate that loss of Her9 causes a significant decrease in rod photoreceptors, subsets of cone photoreceptors, Müller glia cells, and CMZ size. We also find that *her9* mutant photoreceptors have abnormal outer segments and undergo apoptosis. Finally, we show that *her9* expression in the retina is regulated by RA signaling, but RA responsiveness is not completely lost in *her9* mutants. Taken together, our study confirms the requirement for Her9/HES4 in retinal stem cell proliferation and identifies Her9/HES4 as a novel regulator of photoreceptor specification, maintenance, and survival.

## Results

### *Her9* expression during retinal development

Previous studies in zebrafish have described *her9* expression patterns in the early embryo (Bae et al., 2005; Latimer et al.,2005; Leve et al., 2001), and *her9* expression has also been documented in the CMZ of the post-embryonic medaka retina (Reinhardt et al., 2015). We investigated the expression of *her9* during the critical stages of retina development. We used fluorescent in situ hybridization (FISH) to detect *her9* mRNA in the developing zebrafish retina between 24 and 72 hpf, a time period which encompasses the progression from a pseudostratified proliferative retinal neuro-epithelium to a fully laminated, functional retina containing all retinal cell types. At 24 hpf, *her9* was expressed throughout the lens and retina, but most predominately in the ventral portion of the retina, and in a ring around the lens (Fig. 1A-A”). At 48 hpf, *her9* expression became restricted to the undifferentiated peripheral retina, with some expression remaining in the lens (Fig. 1B-B”). By 72 hpf, *her9* expression was only detected in a small patch of cells within the persistently neurogenic CMZ (Fig. 1C-D”). Taken together, the expression of *her9* throughout the undifferentiated embryonic retina and in the proliferative CMZ suggests that it functions in retinal progenitor cells and potentially plays a role in retinal stem cell proliferation or maintenance.

**Figure 1.**
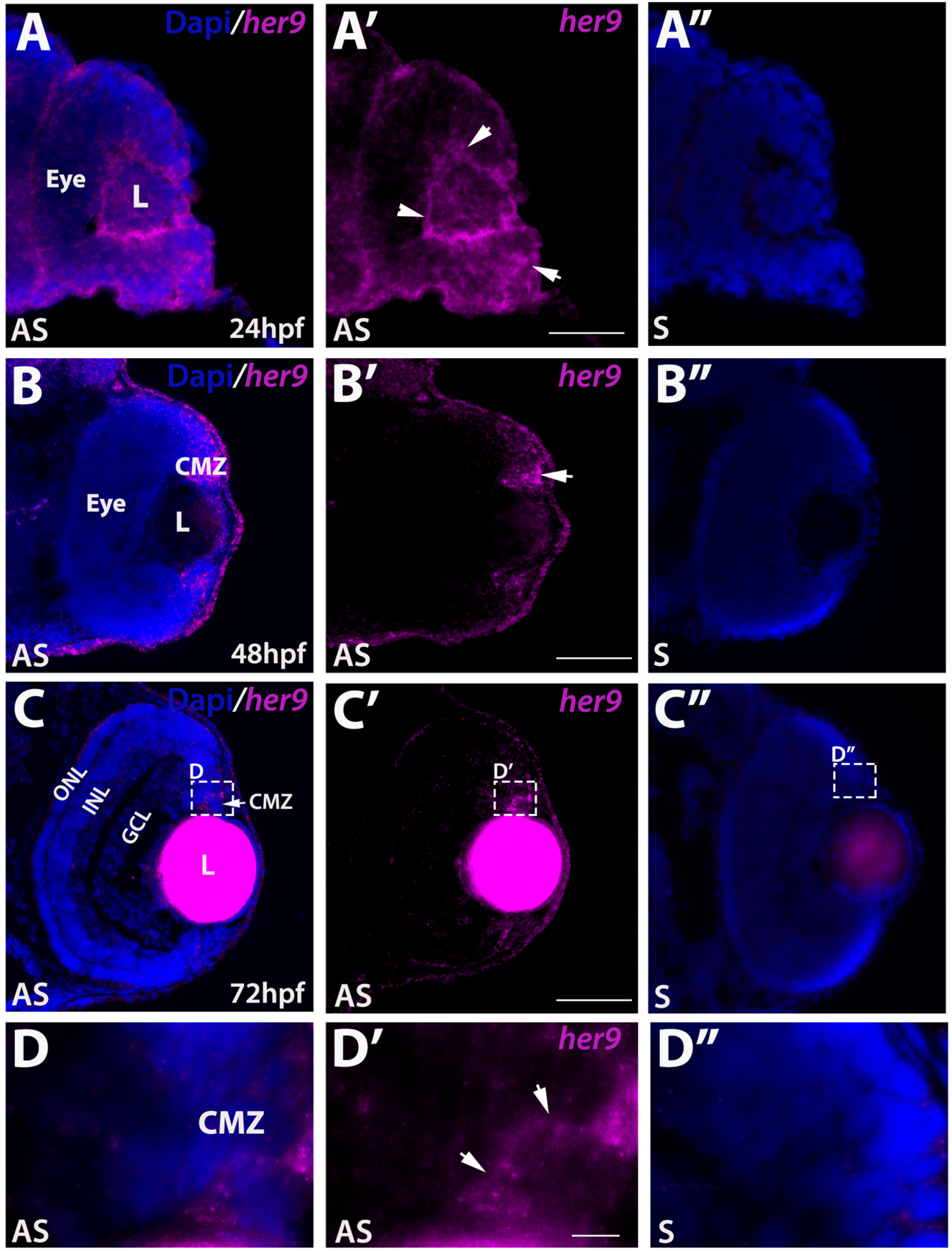
*Her9* expression in the developing retina. Fluorescent in situ hybridization (FISH) showing *her9* mRNA expression at 24 (**A-A’**), 48 (**B-B’**), and 72(**C-C’**) hpf in the developing retina. (**D-D’**) 100x magnification of boxed area in C-C’ showing *her9* expression in the CMZ. *her9* sense probes at 24, 48, and 72 hpf are shown in (**A”, B”, C”, D”**). L, lens; CMZ, ciliary marginal zone; NR, neural retina; ONL, outer nuclear layer; INL, inner nuclear layer; GCL, ganglion cell layer. Scale bar = 50 μm and 100 μm.

### Generating *her9* mutants using CRISPR/Cas9

To explore the role of *her9* during retinal development, we used the CRISPR/Cas9 system to introduce mutations in the *her9* gene, targeting a region of exon 1 upstream of the bHLH domain. We recovered two *her9* mutant alleles, one carrying a 1-bp deletion and the other a 1-bp insertion at the target site (Fig. S1A). Both mutations cause a frameshift resulting in a premature stop codon. Both mutations introduced novel restriction sites, allowing for genotyping by RFLP analysis (Fig. S1B). We used qPCR to investigate *her9* expression in *her9*^*-/-*^ zebrafish, and observed a significant reduction in *her9* mRNA levels, suggesting that the mutations resulted in nonsense-mediated decay (Fig. S1C). We also used a HES4 antibody to detect Her9 protein, which confirmed a significant decrease in expression in *her9* mutants (Fig. S1D). These results indicate that the CRISPR-generated genetic lesion in *her9* is a null mutation. Both mutant alleles presented similar phenotypes so here we present data collected from the 1bp insertion mutant.

### Characterization of the *her9* mutant phenotype

We first characterized the progeny of *her9* heterozygous incrosses by light microscopy. At 24 hpf, *her9* mutants appeared somewhat developmentally delayed with lighter pigmentation and a smaller body size when compared to wild type (WT) and heterozygous siblings (Fig. 2B-C). The mutants also displayed delayed development of the midbrain and hindbrain ventricles (Fig. 2B), which appeared around 36 hpf (data not shown). At 48 hpf, mutant embryos were smaller in body size, microphthalmic, and some displayed pericardial edema (Fig. 2D-E). At 72 hpf, homozygous mutants displayed a phenotypic range from mild to severe, where some mutant embryos looked fairly normal, and others displayed the same defects seen at earlier time points (Fig. 2F-G). In wild type (WT) and *her9* heterozygotes, the swim bladder was fully developed by 5 dpf (Fig. 2A). In contrast, *her9* homozygous mutant larvae did not develop a swim bladder and displayed an abnormal swimming pattern (Fig. 2A, Fig. S2). Other mutant phenotypes observed at 5 dpf included an enlarged liver, curved body, and craniofacial defects (Fig. 2A). At 12 days post fertilization (dpf) we noticed a significant decrease in larval survival with approximately 29% mortality. None of the *her9*^*-/-*^ larvae survived beyond 13 dpf, indicating that loss of Her9 results in larval lethality (Fig. 2H).

**Figure 2.**
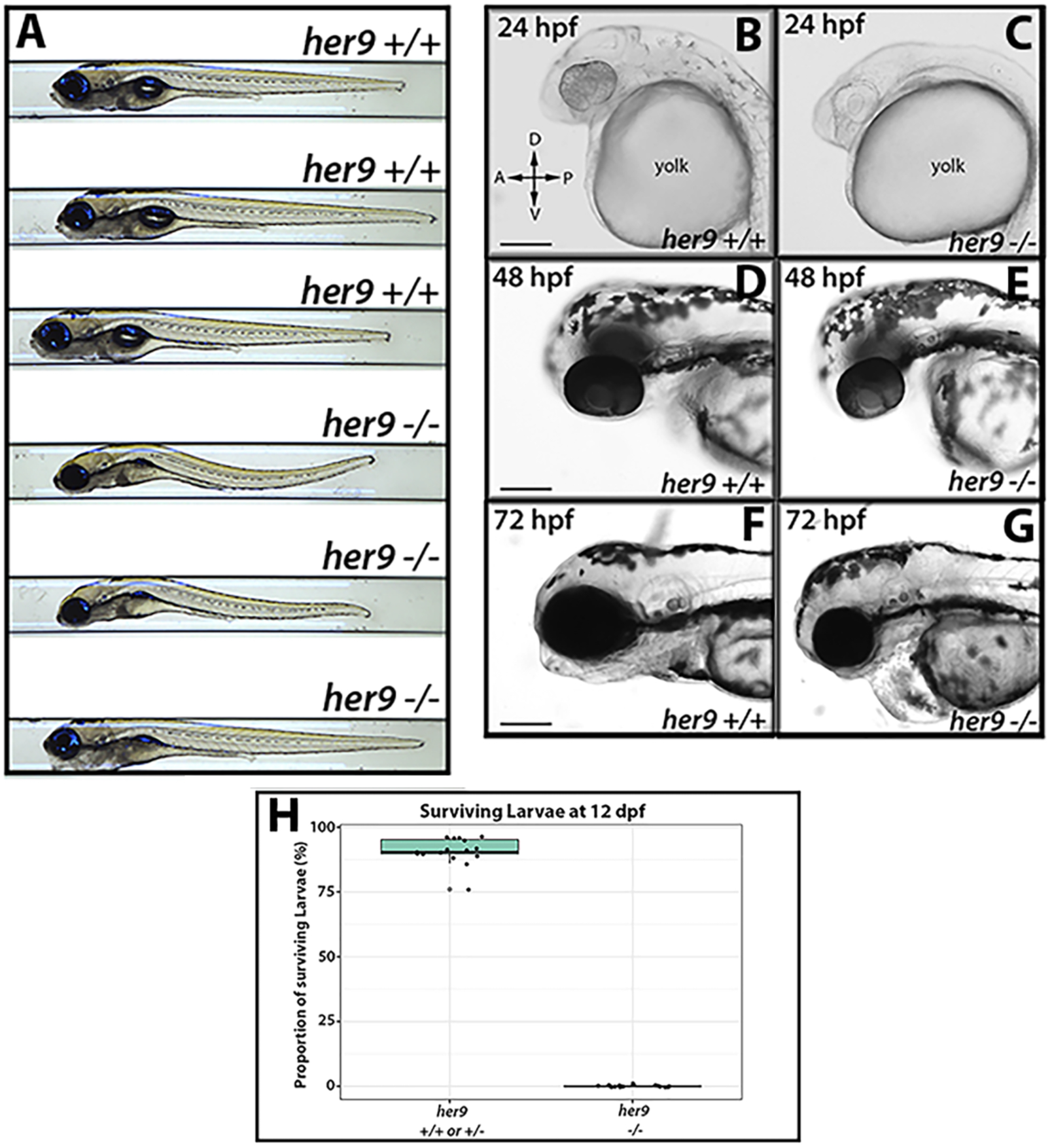
*Her9* mutant phenotype. **(A)** Gross morphology of WT and *her9* mutants at 5 dpf. (**B-G)** Gross morphology of WT and *her9* mutants at 24, 48, and 72 hpf. (**H)** Proportion of surviving larvae at 12 dpf; each point represents a separate cross. Scale bar= 50 µm and 100 µm.

To rule out off-target contributions to the mutant phenotype, we performed a *her9* mRNA rescue experiment. Full-length *her9* mRNA was microinjected into one-cell stage embryos from *her9* heterozygous in-crosses, then the embryos were imaged and scored for the earliest visible mutant phenotype, which was lack of a clearly defined midbrain-hindbrain boundary and a missing midcerebral vein (MeCV) at 24 hpf (imaged by fluorescence microscopy of the *fli1*:GFP transgene; Fig. S3A-C). Whereas only 5.9% of uninjected *her9* homozygous mutant larvae had a detectable MeCV at 24 hpf, 80% of the *her9* mutant embryos that received *her9* mRNA had a MeCV (Fig. S3C-D). We also observed a significant increase in the vasculature around the eye and heart in mRNA-injected mutant embryos (Fig. S3B-C). As further confirmation that the phenotypes observed are specific to loss of Her9, the 1bp insertion heterozygotes were crossed with the 1bp deletion heterozygotes and the resulting phenotype in compound heterozygotes was equivalent to that seen in the individual mutants (not shown). Combined, these results indicate that the phenotypes described herein are specifically due to the mutation of *her9*, and also that Her9 may be required for proper vasculature development.

### *Her9* mutants lack a visual background adaptation response and display visual dysfunction

We observed that at 5 dpf, a significant number of the progeny of *her9*^*+/-*^ incrosses (∼27%) appeared more darkly pigmented after prolonged light exposure when compared to their siblings. This suggested that *her9* mutants may lack a visually mediated background adaptation (VBA) response, a neuroendocrine camouflage response that allows zebrafish to manipulate their melanin granules in response to light exposure to match their background. Several previous studies have shown that the VBA response depends on retinal input (Mueller and Neuhauss, 2014; Neuhauss et al., 1999; Viringipurampeer et al., 2014), thus an absent VBA response can indicate visual impairment.

To determine whether the lack of a VBA response was associated with the *her9* mutant genotype, we first dark-adapted the larvae for two hours, then placed them in ambient light for 30 minutes, imaged and scored them as dark or light, then extracted genomic DNA for genotyping. After light exposure, 51 out of 70 larvae were scored as light and 100% of those embryos genotyped as WT or heterozygous (p< 0.0001; Fig. 3A and C). In contrast, 19 of the larvae scored as dark, 63% of which genotyped as homozygous mutants (p= 0.0051; Fig. 3B-C). From these data, we conclude that *her9* mutants lack a normal VBA response, which could suggest visual impairment and possible retinal defects.

**Figure 3.**
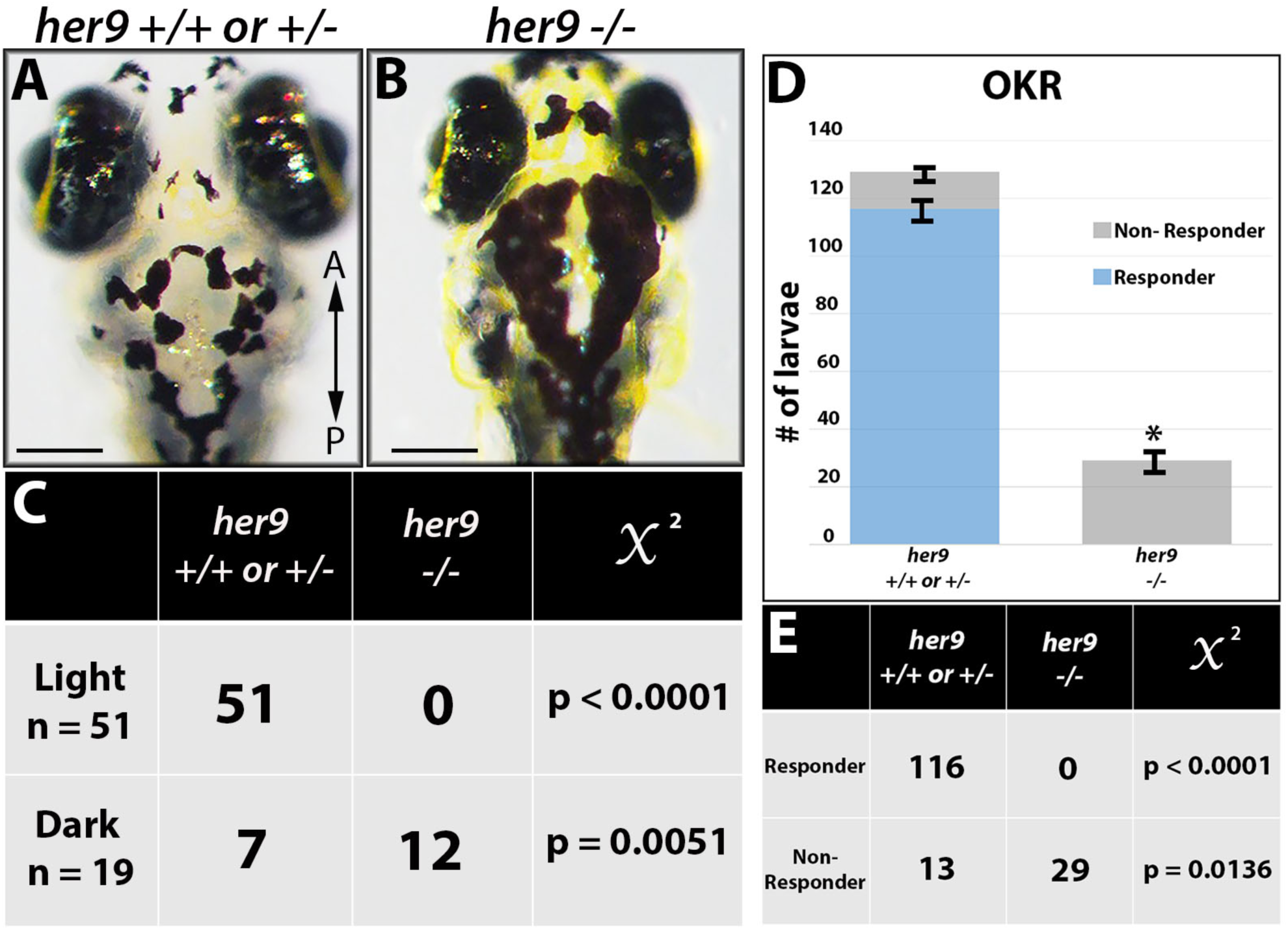
Her9 mutants lack a VBA response and display abnormal visual behavior (OKR). **(A-B)** Images displaying *her9* mutant with significantly darker pigmentation at 5 dpf **(C)** Genotyping of the “dark” and “light” larvae revealed that out of 51 individuals classified as “light”, zero were *her9*^*-*/-^ (p-value<0.0001, Chi-square analysis). Conversely, 12 out of 19 individuals classified as “dark” were *her9*^*-*/-^ (p-value=0.0051). (**D**) Number of responders and non-responders in the optokinetic response (OKR) assay. **(E)** Genotyping after OKR revealed that zero of the *her9*^*-/-*^ embryos displayed saccades in the OKR (p<0.0001). Scale bar= 50 µm.

Given their lack of VBA response, we wondered whether the *her9* mutants had any functional visual deficit. To investigate this, we used the optokinetic response (OKR) assay, which is a behavioral test that measures the larvae’s ability to perform a combination of smooth pursuit and rapid saccade eye movements in response to a moving pattern of alternating black and white vertical stripes (Brockerhoff, 2006). Individual larvae from a *her9* heterozygous incross were screened at 5 dpf over three 30-sec trials (n= 159), and then genotyped after screening. Whereas 116 out of 129 WT and heterozygous larvae displayed a positive OKR response, none of the *her9* mutants responded with saccadic movements in the OKR assay (0 out of 29 tested; p<0.0001; Fig. 3D-E). We conclude from these data that the loss of Her9 causes impaired visual responses at 5 dpf.

### *Her9* mutants display a decrease in rod photoreceptors and rod outer segment defects

Next, we used immunohistochemistry (IHC) on retinal tissue sections to examine the number and morphology of various retinal cell types in WT and *her9* mutant larvae, starting with the rod photoreceptors. Using a rod photoreceptor specific antibody (4C12), we observed numerous rod photoreceptors in the dorsal and ventral portions of the WT retina at 12 dpf (Fig. 4A). At higher magnification the rod outer segments were easily detected (Fig. 4A’). In contrast, *her9* mutant retinas displayed a significant decrease in rod photoreceptors which was especially apparent in the dorsal retina (Fig. 4B). At higher magnification we also noticed that the outer segments of the remaining *her9* mutant rods looked shorter and distorted compared to those in the WT retina.

**Figure 4.**
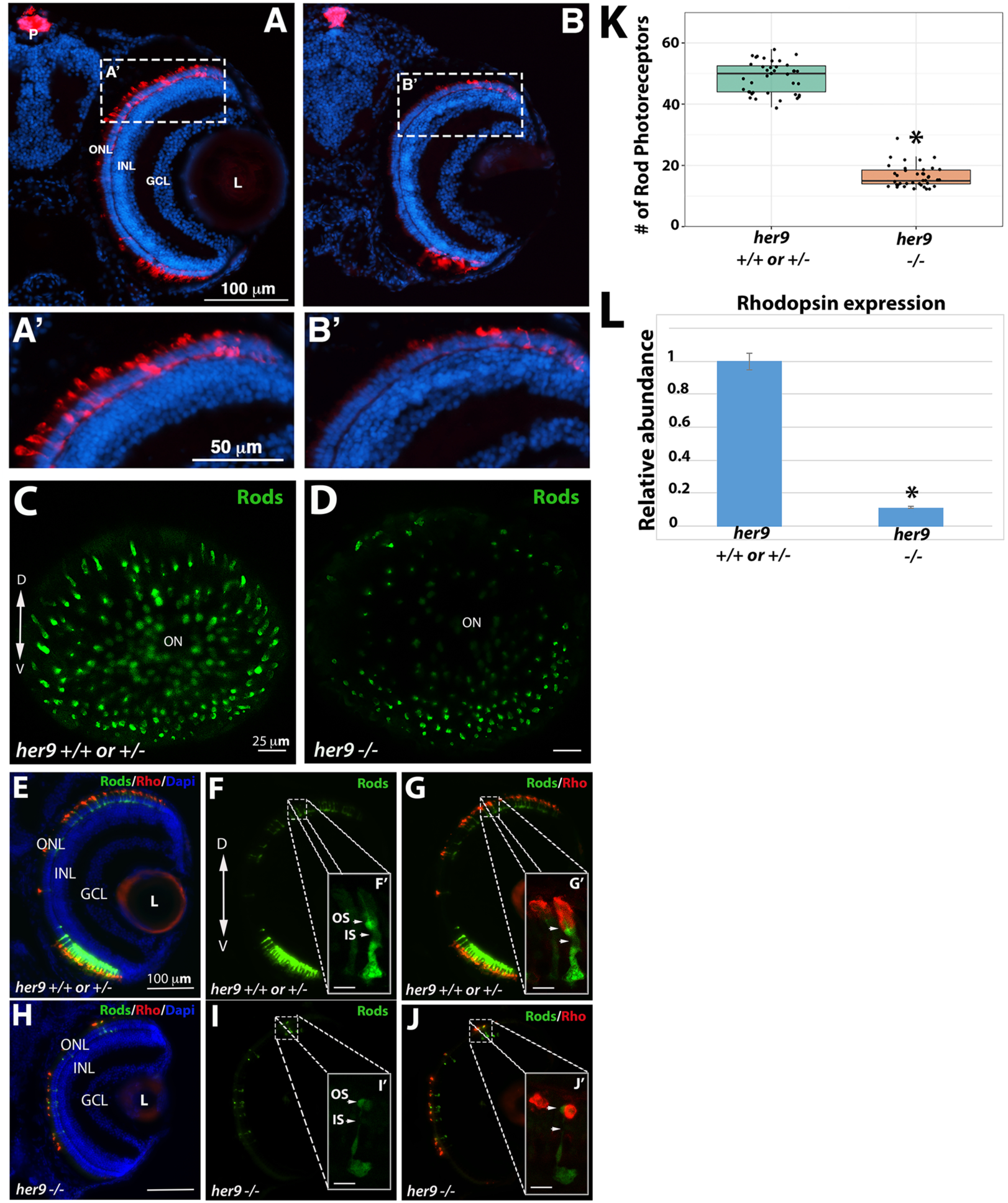
*Her9* mutants have fewer rods with shorter outer segments. Immunohistochemistry with a rod antibody (4C12) in *her9*^+/+ or +/-^ (**A, A’**) and *her9*^-/-^ (**B, B’**) retinal sections. **(C-D)** Confocal images of whole eyes from *her9* heterozygous incross progeny on the XOPS:GFP background. **(E-G)** Immunohistochemistry with an antibody that labels rhodopsin (1D1) on retinal cryosections of XOPS:GFP WT and heterozygous **(E-G)** or *her9* mutant (**H-J**) larvae. **(K)** Rod cell counts in *her9* ^*-/-*^ larvae and their WT and heterozygous siblings. **(L)** qPCR analysis of *rhodopsin* expression at 8 dpf (fold change relative to *Ef1α*). ONL, outer nuclear layer; OPL, outer plexiform layer; INL, inner nuclear layer; GCL, ganglion cell layer; L, lens; ON, optic nerve; P, pineal gland. Scale bar= 50 µm and 100 µm.

To more thoroughly examine the effects of the *her9* mutation on rod photoreceptor number and morphology, *her9* heterozygotes were crossed onto the XOPs:GFP transgenic background, which fluorescently labels rod photoreceptors (Fadool, 2003). The *her9* heterozygotes on the XOPs:GFP background were in-crossed, and whole eyes were removed from larvae at 5 dpf and imaged by confocal microscopy, while the rest of the body was used for genotyping. Using this approach, we observed a significant decrease in GFP^+^ rod photoreceptors across the entire *her9* mutant retina when compared to the WT (Fig. 4C-D), confirming our previous results.

The remaining heads were pooled together based on genotype and cryo-sectioned, followed by counting of GFP+ cells. We observed a 69% reduction in the number of GFP+ rods in *her9* mutant larvae compared to WT (avg. 18 per 100 μm in *her9* mutants vs. 58 per 100 μm in WT), which was confirmed via qPCR for *rhodopsin* (Fig. 4K-L). Finally, we used the 1D1 antibody to label the rod outer segments in WT and *her9* mutant retinal cryosections. In the *her9* mutant embryos the rod outer segments appeared severely truncated compared to the outer segments of the WT embryos (Fig. 4E-G’ and 4H-J’). Taken together, these data demonstrate that loss of Her9 causes a large decrease in rod photoreceptor number and defects in rod photoreceptor outer segment morphology.

### Cone outer segments are truncated in *her9* mutants

We used the Zpr1 antibody to detect the red-green double cones in WT and *her9* mutant retinas. At 12 dpf, we initially did not observe an obvious decrease in cone number in the *her9* mutant retina (Fig. 5A-B). However, at higher magnification, we detected gaps in the spacing of the double cones, as well as truncated cone outer segments, in *her9* mutant retinas (Fig. 5A’-B’). To examine this further, we crossed the *her9* mutation onto the TαC:GFP transgenic background, which fluorescently labels all cone photoreceptor subtypes (Kennedy et al., 2001;Fig. 5C-F). There was a significant decrease in the number of cone photoreceptors in the *her9* mutants compared to WT (Fig. 5I). We used the Zpr3 antibody to label the outer segments of the double cones and rods, and found that these were missing or truncated in *her9* mutant retinas (Fig. 5E-E’ and 5H-H’). These results demonstrate that the loss of Her9 causes a decrease in the number of double cones, and has profound effects on red and green cone outer segment morphology.

**Figure 5.**
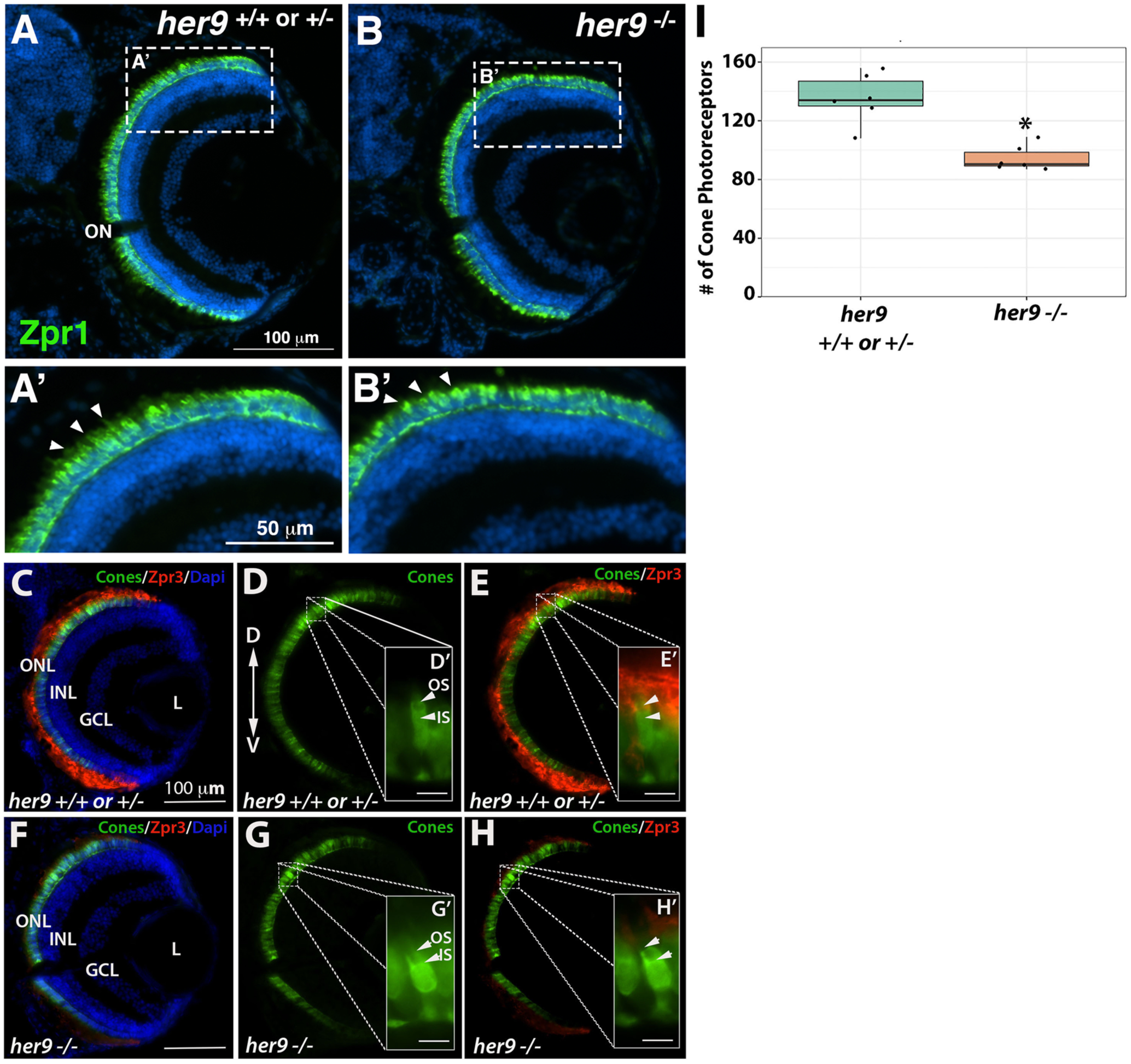
Cone outer segments are truncated in *her9* mutants. Immunohistochemistry with a red-green cone antibody (Zpr1) in *her9*^+/+ or +/-^ (**A, A’**) and *her9*^-/-^ (**B, B’**) retinal sections. Arrowheads indicate outer segments. **(C-H)** Immunohistochemistry with an antibody that labels rod and double cone outer segments (Zpr3) on retinal cryosections of T***α***C:GFP;*her9*^*+/+ or +/-*^ (**C-E**) or T***α***C:GFP;*her9*^*-/-*^ mutant (**F-H**) larvae. **(I)** Cone cell counts in *her9*^*-/-*^ mutant larvae and their WT and heterozygous siblings. ONL, outer nuclear layer; OPL, outer plexiform layer; INL, inner nuclear layer; GCL, ganglion cell layer; L, lens; ON, optic nerve; OS, outer segment; IS, inner segment. Scale bar= 100 µm

### *Her9* mutants display cone subtype-specific phenotypes

We next investigated whether all cone subtypes were equally affected by the loss of Her9, using cone opsin-specific antibodies to perform IHCs on retinal cryosections as described above. We again observed a significant reduction (42%) in the number of green cone photoreceptors and truncated green cone outer segments in the *her9* mutant retinas compared to their WT siblings (Fig. 6A-B, G). Interestingly, IHC with the UV- and blue-cone opsin antibodies revealed a much smaller decrease in number for those cone subtypes in *her9* mutant retinas compared to the decrease in green cones (6.6% and 5%, respectively; Fig. 6C-F’, H and I) and we did not detect truncation in the blue and UV cone outer segments (Fig. 6D’-F’). qPCR for long-, medium- and short-wavelength opsins confirmed a small decrease in UV- and blue-cone opsin expression and a large decrease in red cone opsin expression (Fig. 6J). Green cone opsin expression was also modestly reduced by qPCR. The results of these experiments suggest that loss of Her9 causes variable reductions in all cone photoreceptors and defects in outer segments that are specific to red/green cones and rods.

**Figure 6.**
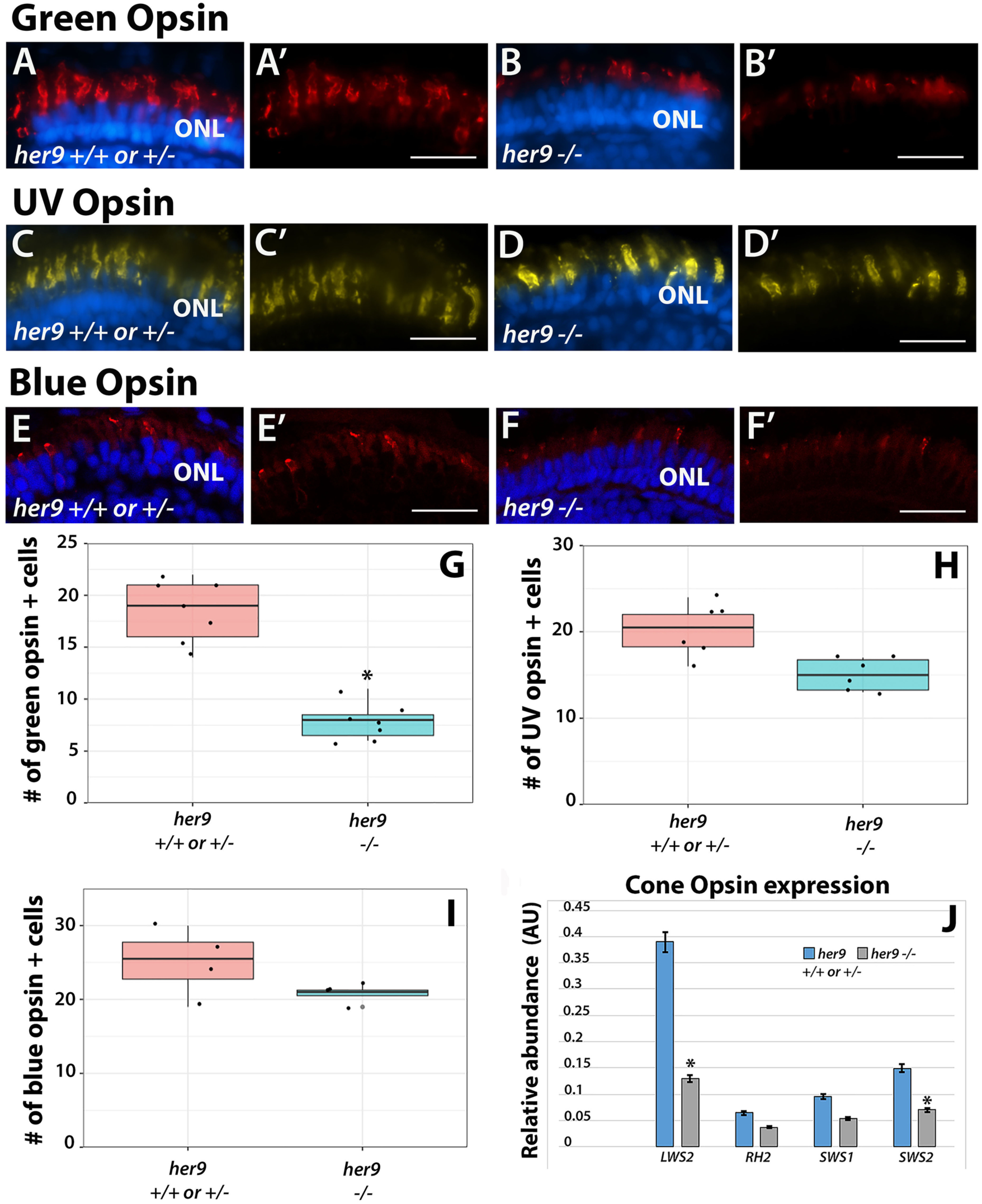
*Her9* mutants display cone subtype-specific phenotypes. Immunohistochemistry using a green **(A-B’)**, UV **(C-D’)**, or blue **(E-F’)** cone opsin antibody on WT and *her9* mutant retinal cryosections. **(G-I)** Cell counts of opsin expressing cells in *her9* mutant retinas compared to their siblings (normalized to eye size). **(J)** qPCR analysis of the different cone opsins at 8 dpf (fold change relative to *Ef1α*). ONL, outer nuclear layer. Scale bar = 50 µm.

### Müller glia abnormalities in *her9* mutants

To determine whether loss of Her9 affects other late-born cell types, we crossed the *her9* mutation onto the gfap:GFP transgenic background, which fluorescently labels retinal Müller glia (Li et al., 2015). At 5 dpf, *her9* homozygous mutants showed a significant reduction of GFP+ Müller glia compared to their WT and heterozygous siblings (p<0.0001; Fig. 7A-C). We also observed a decrease in GFP+ glial cells across the entire *her9* mutant embryo (brain and enteric nervous system) at this stage (not shown). In addition, the GFP+ Müller glia that were present in *her9* mutant retinas displayed a disorganized pattern, with their cell bodies located at various depths of the INL layer, rather than forming a uniform row as in the WT retinas (Fig. 7A’ and B’). We also observed a lack of processes from the Müller glia spanning the retina in the *her9* mutants compared to their WT siblings. These data indicate that the loss of Her9 not only causes a reduction in the number of Müller glia, but also distorts the morphology, organization and patterning of the Müller glial cells that do develop.

**Figure 7.**
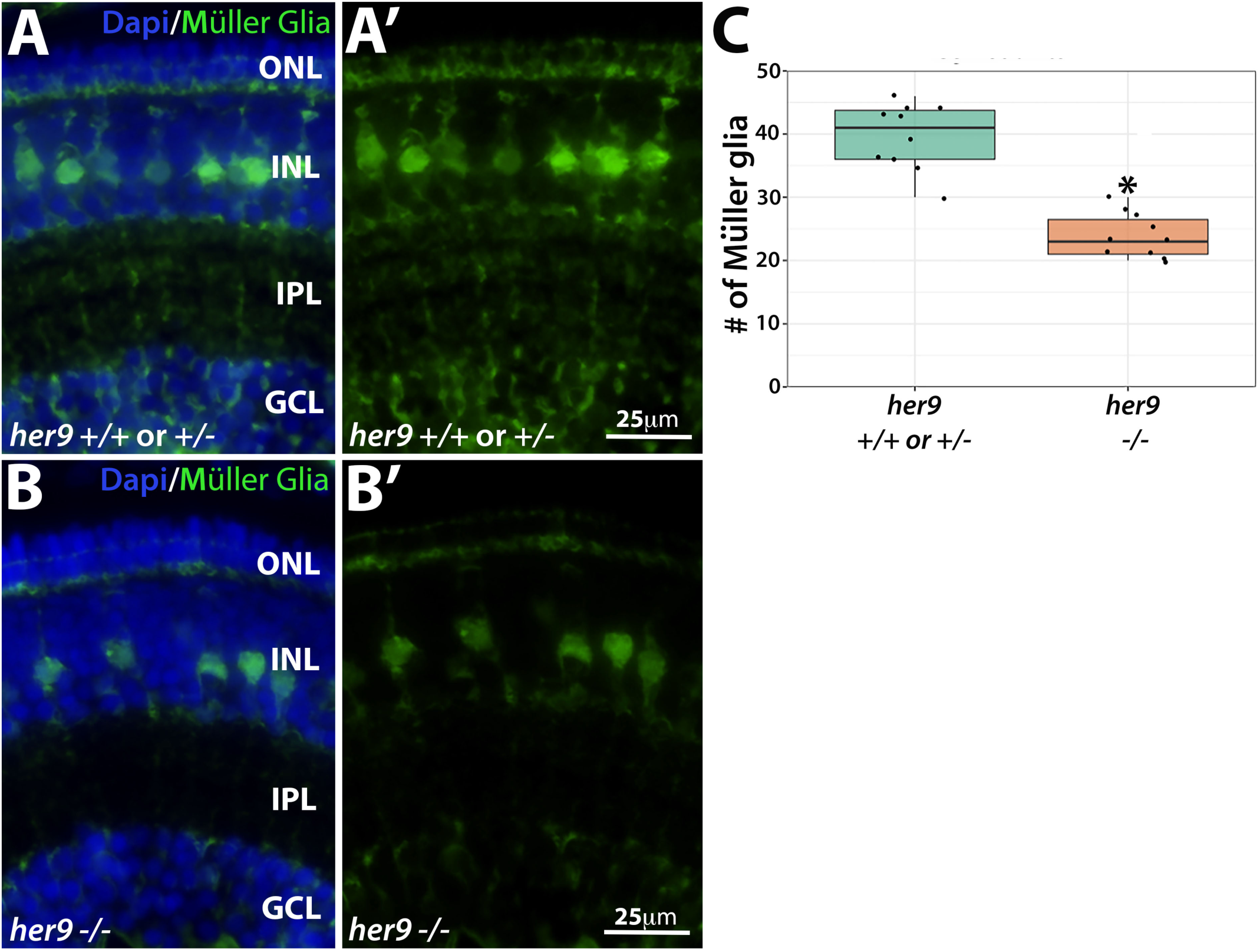
Loss of *her9* causes a decrease in Müller glia number and distorts their organization. Cryosections of WT and heterozygous **(A-A’) or** *her9* mutant **(B-B’)** retinas at 5 dpf on the gfap:GFP transgenic background. (**C**) Cell counts of GFP+ Müller glial cells in *her9* mutant retinas compared to their WT siblings (normalized to eye size).

### Loss of Her9 has minimal effects on other retinal cell types

Does the loss of Her9 disrupt development of all retinal cell types? To address this question, we used transgenic lines or IHC with cell-type specific antibodies for ganglion cells, amacrine cells, and horizontal cells. First, we crossed *her9* heterozygous zebrafish onto the ath5:GFP transgenic background (Masai et al., 2003) which fluorescently labels ganglion cells and the optic nerve (Fig. 8A-B). We observed a modest decrease in GFP expression in the *her9* mutant retinas compared to WT and somewhat thinner optic nerves (Fig. 8B; arrow). We followed this up with IHC using the HuC/D antibody, which labels both ganglion cells and amacrine cells. We observed a significant decrease in the number of ganglion cells in *her9* mutant retinas compared to the WT and heterozygous siblings (Fig. 8C-F, G; p < 0.0001). However, the number of amacrine cells was not significantly different in the WT versus *her9* mutant retinas at 5 dpf (not shown).

**Figure 8.**
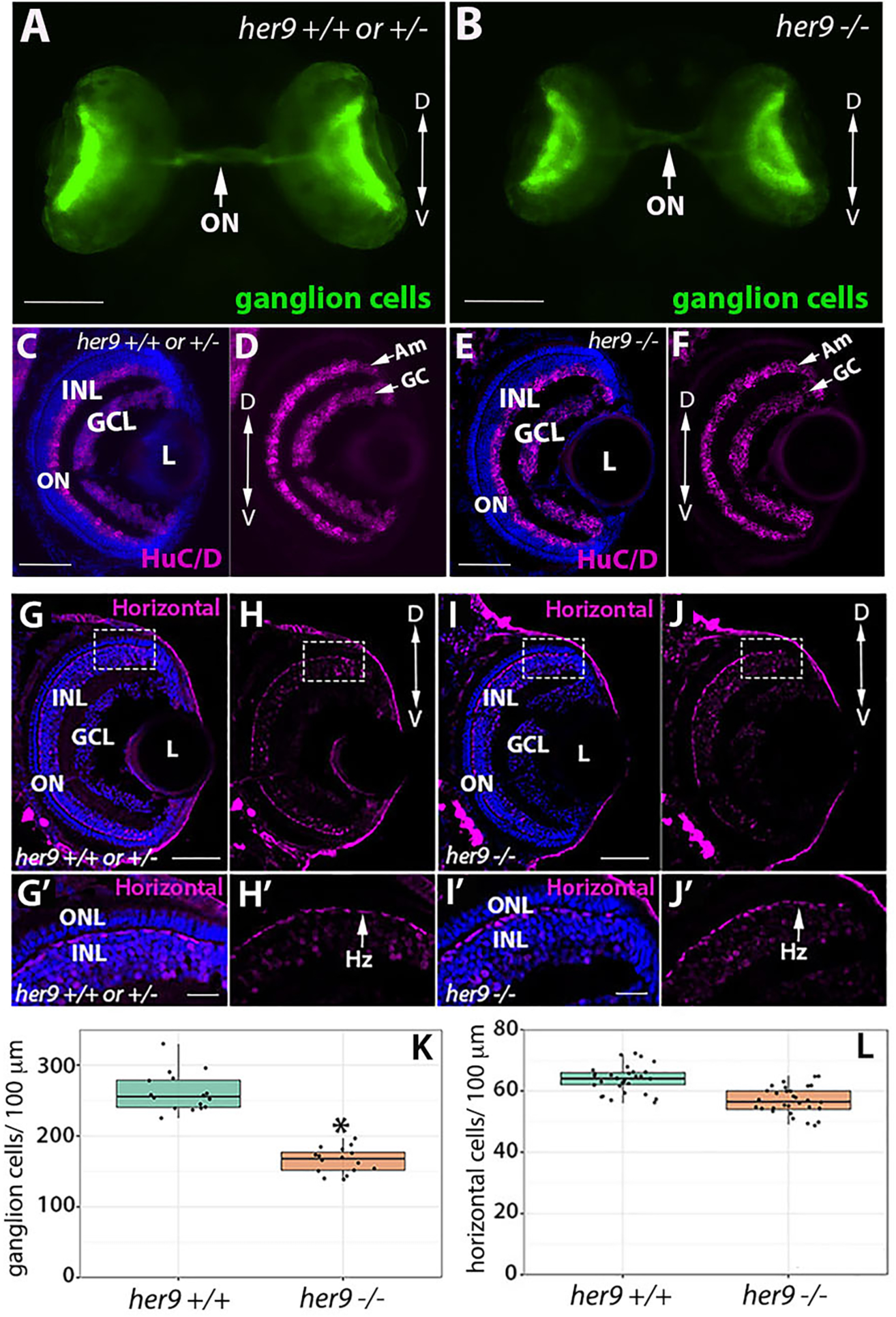
Loss of *her9* has minimal effects other retinal cell types. Whole eye images of WT and heterozygous embryos **(A)** or *her9* mutant embryos **(B)** on the Ath5:GFP transgenic background at 5 dpf. Immunohistochemistry for ganglion and amacrine cells (HuC/D antibody) in cryosections from WT and heterozygous retinas **(C-** or *her9* mutant retinas **(E-F). (G-H’)** Immunohistochemistry with the Prox1 antibody labels horizontal cells in cryosections of WT and heterozygous (**G-H’**) or *her9* mutant **(I-J’)** retinas. Cell counts reveal a decrease in the number of ganglion cells in the mutant retinas (**K**) but no significant difference in the number of horizontal cells (**L**). Am, Amacrine cells; GC, Ganglion cells; Hz, Horizontal cells; ONL, outer nuclear layer; INL, inner nuclear layer; ON, optic nerve; L, Lens. Scale bar= 50 µm and 100 µm.

The Prox1 antibody was then used to assess horizontal cells in the retinas of *her9+/-* in-cross progeny (Fig. 8G-J’). There were no morphological differences in the horizontal cells of *her9* mutants compared to WT (Fig. 8H’ and J’). A small decrease in the number of horizontal cells was observed in mutant retinas compared to WT, but it was not statistically significant when normalized to the smaller mutant eye size (Fig. 8L). These results demonstrate that loss of Her9 causes a mild decrease in ganglion cell number, but does not alter the numbers of amacrine or horizontal cells, and does not disrupt the morphology of these retinal cell types. This indicates that the morphological defects due to loss of Her9 are specific to photoreceptors and the Müller glia.

### Loss of Her9 causes a progressive collapse of the CMZ

Previous studies of the post-embryonic Medaka and Xenopus retina showed that *her9* regulates the proliferation of the retinal stem cells in the CMZ (Reinhardt et al., 2015). Therefore, we wanted to investigate whether germline mutation of zebrafish *her9* would affect the establishment of the CMZ during retinal development. We collected zebrafish larvae at 72 hpf, then prepared retinal sections for PCNA immunolabeling, which detects cells in S-phase (Fig. 9A-A’). We observed a significant decrease in the number of PCNA+ cells in the CMZ of mutant retinas in comparison to their WT and heterozygous siblings. To determine whether the *her9* mutant CMZ could recover to a normal size later in development, we immunolabeled 5 dpf retinal sections with the PCNA antibody. We again observed significantly fewer PCNA+ cells in the CMZ of 5 dpf *her9* mutant retinas compared to their WT and heterozygous siblings (Fig. 9B-B’). Quantification of both the PCNA-labeled CMZ area as well as fluorescence intensity confirmed a significant decrease in the *her9* mutants compared to WT and heterozygous siblings (Fig. 9C-D). Interestingly, the lack of proliferating cells in the CMZ became more severe from 72 hpf to 5 dpf in *her9* mutants, indicating that there is a decrease in the number of proliferating cells in the mutant CMZ over time. Our results confirm that Her9 regulates the proliferation of stem cells in the CMZ. Moreover, as we also observed a decrease in pockets of stem cells in the *her9* mutant brain (not shown), this suggests that Her9 has a general role in regulating stem cell proliferation or maintenance throughout the developing central nervous system.

**Figure 9.**
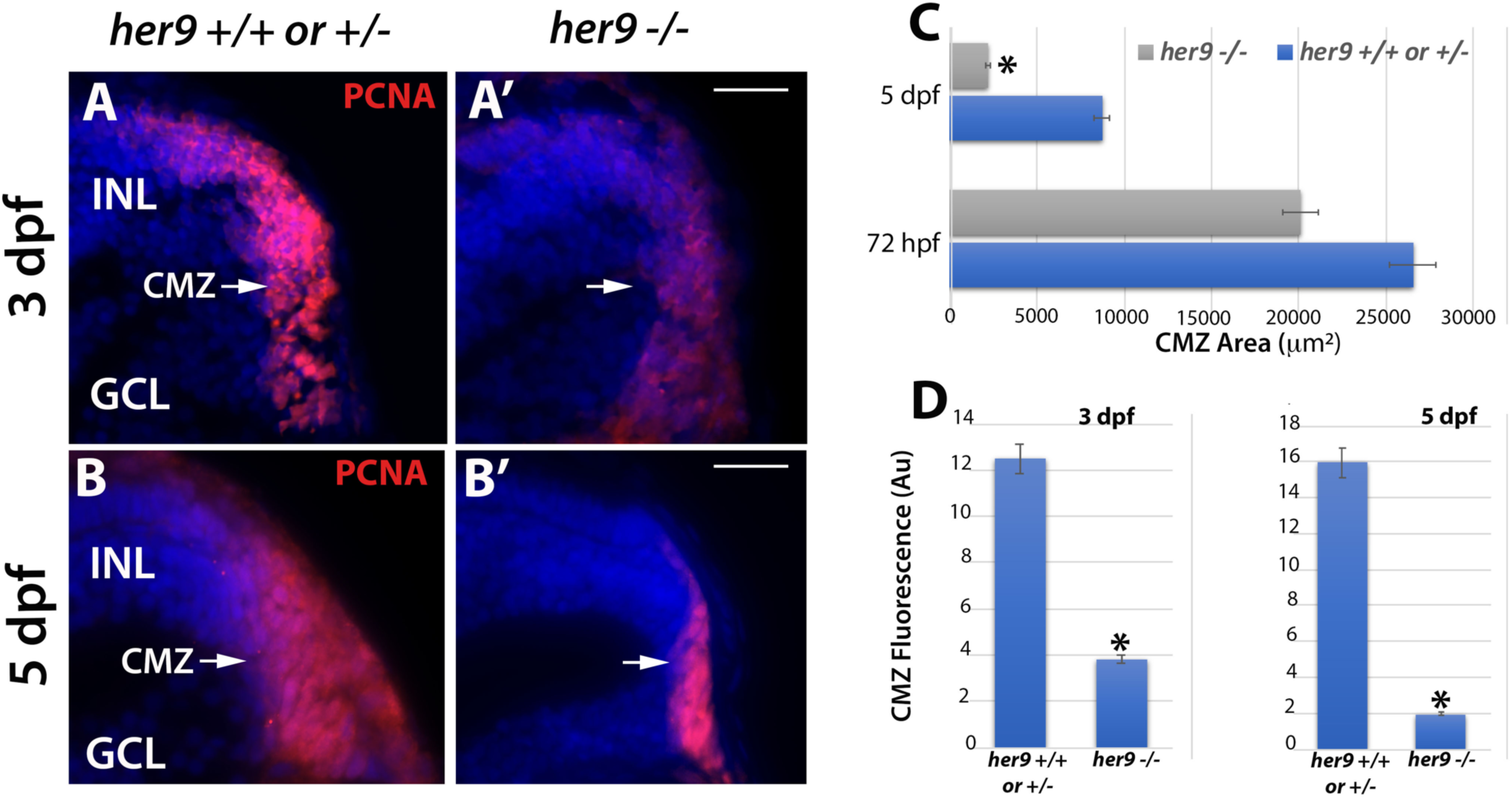
CMZ defects in *her9* mutants. **Immunostaining for** PCNA in WT or heterozygous (**A, B**) and *her9* mutant (**A’, B’**) retinal sections at 72 hpf and 5 dpf; a decrease in the size and fluorescence intensity of the CMZ is apparent in *her9* mutants. Area of CMZ calculated at 72 hpf and 5 dpf (**C**). *Her9* mutants have significantly fewer PCNA-positive cells in the peripheral retina. CMZ fluorescence intensity at 72 hpf and 5 dpf (**D**). GCL, ganglion cell layer; INL, inner nuclear layer; CMZ, ciliary marginal zone. Scale bar = 50 µm.

### *Her9* mutants display abnormal expression of photoreceptor lineage genes

To begin to address the mechanism underlying the photoreceptor phenotypes of *her9* mutant retinas, we first examined whether loss of *her9* affects the specification of retinal progenitor cells towards rod and cone lineages. Crx is a transcription factor that plays a critical role in the specification of all photoreceptor subtypes (Rath et al., 2007). We used FISH on retinal cryosections to examine the expression of *crx* at 48 and 72 hpf. Starting at 48 hpf, the expression of *crx* spreads across the WT retina in a ventral to dorsal fan-like manner (Fig. 10A-A’). In *her9* mutant retinas we observed a similar expression pattern of *crx*, albeit with reduced signal intensity (Fig. 10B-B’); by 72 hpf, *crx* expression in *her9* mutant retinas was not significantly different from WT (Supplemental Fig. S4A-B’). Therefore, we conclude that loss of Her9 does not perturb specification of photoreceptor progenitors.

**Figure 10.**
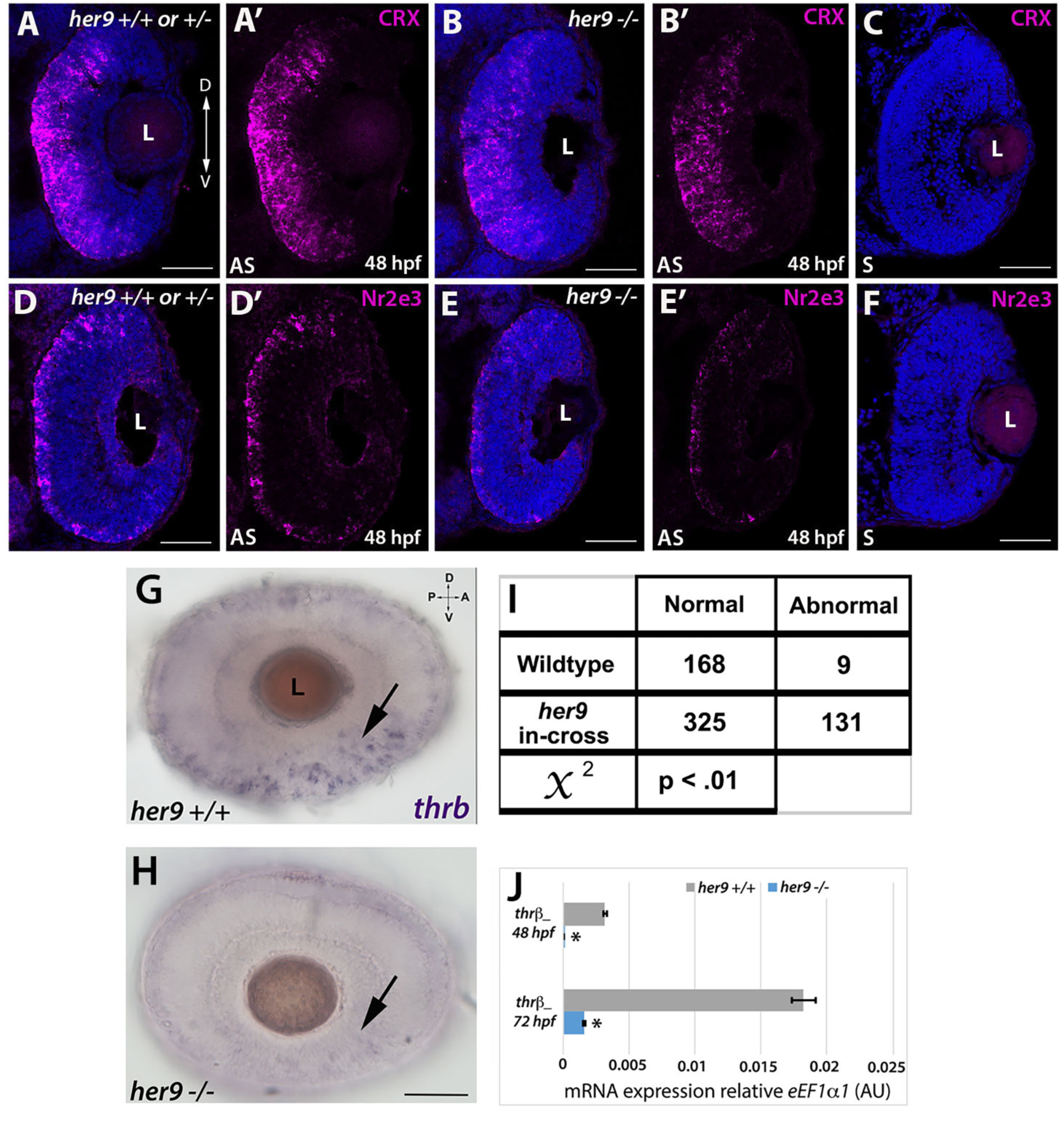
*Her9* mutants display abnormal expression of photoreceptor lineage genes. (**A-B’)** Fluorescent in situ hybridization (FISH) for *crx* at 48 hpf in WT and mutant neural retina. **(C)** *Crx* sense probe. **(D-E’)** Decreased expression of *Nr2e3* in the mutant retina compared to the wildtype. **(F)** *Nr2e3* sense probe. **(G-H).** Whole mount in situ hybridization (WISH) for *thrβ* expression at 48 hpf in WT and *her9* mutant retina. **(I)** Chi-square analysis comparing WT incross and *her9* heterozygous incross expression pattern. **(J)** qPCR for *thr*β at 48 and 72 hpf. L, lens. Scale bar= 50µm

Next, we examined expression of photoreceptor subtype-specific transcription factors. Nr2e3 is responsible for the activation of rod photoreceptor genes and repression of cone genes in photoreceptor progenitors (Chen et al., 2005). At 48 hpf in the WT retina, we observed robust expression of *Nr2e3* throughout the ONL of the retina (Fig. 10D-D’). In contrast, in *her9* mutants, there was significantly less *Nr2e3* expression within the ONL in comparison to the WT (Fig. 10E-E’). At 72 hpf the expression of *Nr2e3* in the *her9* mutant retina had increased relative to 48 hpf, but the expression pattern remained disorganized relative to WT (Fig. S4C-D’). Taken together, this result indicates that the rod photoreceptors in *her9* mutants are specified but their differentiation is abnormal.

Next, we investigated whether the red cone photoreceptor lineage was disrupted in *her9* mutants. To do this, we used whole mount in situ hybridization (WISH) to investigate the expression of *trβ2* in the developing retina at 48 and 72 hpf in progeny from WT and *her9* heterozygous incrosses. The eyes were then removed from the embryos for imaging by light microscopy. In WT larvae, most of the individuals displayed robust expression of *trβ2* in the ventral portion of the eye and along the ONL circumference where the cone photoreceptors are located (the “normal” pattern), although a few WT larval eyes (5.3%) showed fainter expression in the ventral portion of the eye and in the ONL (n=9 out of 177; Fig. 10G-H). In contrast, 28.7% of the larval eyes from *her9* heterozygous incrosses (n=131 out of 456) displayed an abnormal pattern of *trβ2* expression (*χ*^2^ analysis, p<0.01; Fig. 10I). qPCR analysis at 48 hpf and 72 hpf confirmed the reduced expression of *trβ2* in *her9* mutants compared to their wildtype siblings (Fig. 10J). Therefore, we conclude that loss of Her9 disrupts the specification of the red cone lineage.

### *Her9* mutant photoreceptors undergo apoptosis

In addition to abnormal specification, the reduced numbers of rods and double cones in *her9* mutants could reflect a decrease in survival of differentiated photoreceptors. To determine whether this was the case, we performed TUNEL labeling on retinal cryosections from WT and *her9* mutant retinas from 24 hpf to 5 dpf to identify apoptotic cells. At 24 hpf, we observed a slight increase in TUNEL+ cells in the lens of *her9* mutants compared to WT, but no cell death in the retina (Fig. 11A-A’ and E). At 48 hpf, there was no difference in TUNEL+ cells between WT and *her9* mutant retinas (Fig. 11B-B’ and E). However, at 72 hpf we observed a significant increase in TUNEL+ cells in the GCL, INL, ONL, and CMZ of *her9* mutant retinas compared to the WT (Fig. 11C-C’ and E-F). Interestingly, we observed a noticeable cluster of TUNEL+ cells in the dorsal peripheral retina adjacent to the CMZ (Fig. 11C’). Additionally, some of the TUNEL+ cells in the INL had the morphology of Müller glia, which could indicate the Müller glia were dying, but could also be due to Müller glia phagocytosis of other dying cells (Fig. 11C’). By 5 dpf, apoptosis in *her9* mutants remained elevated in the ONL, CMZ, and INL relative to WT retinas (Fig. 11D-D’ and G). These data indicate that Her9 is required for the survival of post-embryonic retinal progenitor cells, photoreceptors, and possibly cells in the INL.

**Figure 11.**
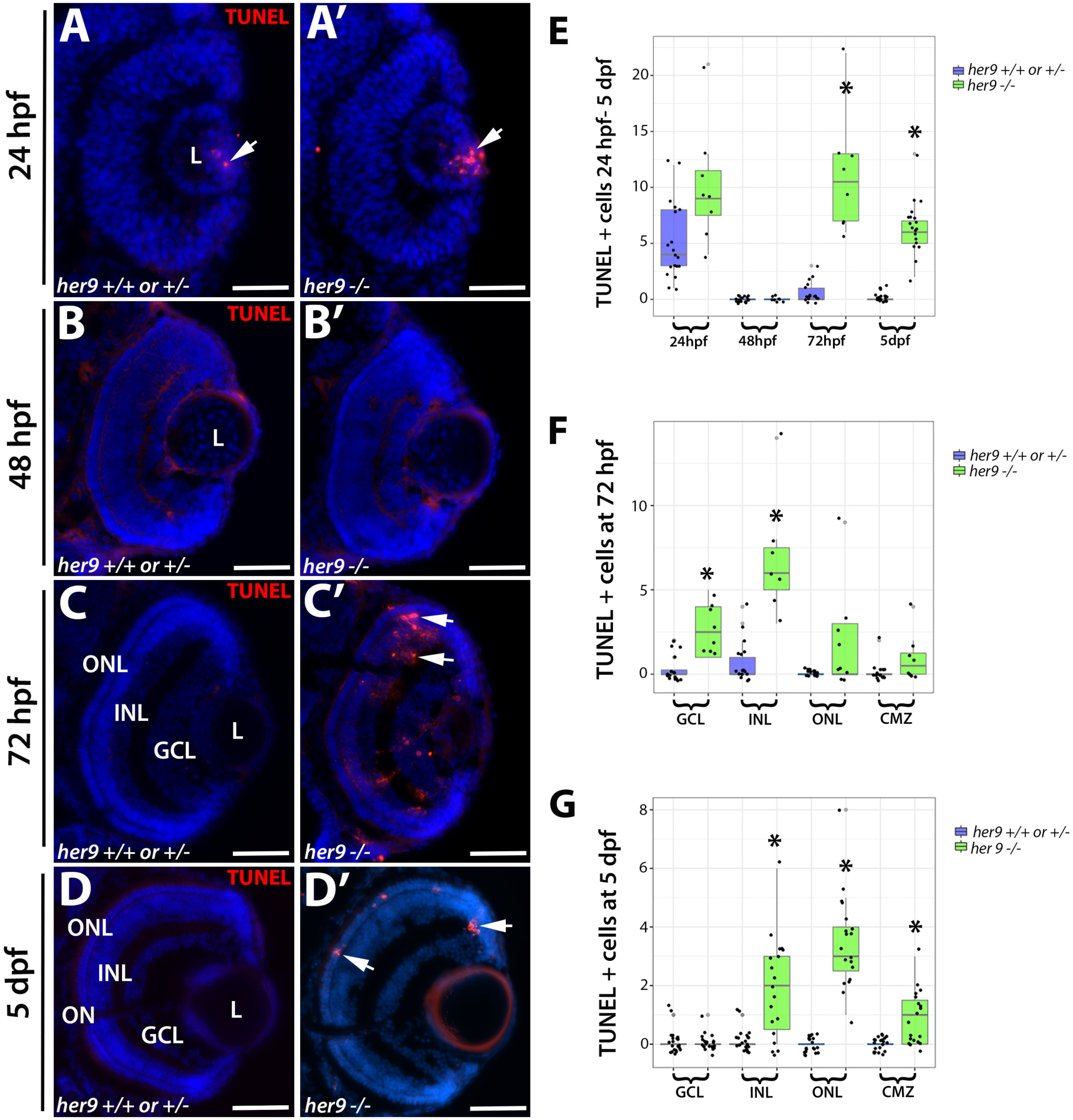
Her9 mutant retinas display increased apoptosis beginning at 72 hpf. **(A-D’)**TUNEL staining on cryosections of WT and *her9* mutant retinas. *Her9* mutant embryos show no significant increase in cell death in the retina at 24 and 48 hpf compared to WT siblings **(A-B’).** At 72hpf, *her9* mutant larvae display increased apoptosis in the GCL, INL, ONL and CMZ (C-C’) that remains elevated in the ONL at 5 dpf (**D-D’**). Cell counts of TUNEL+ cells (**E-G**). ONL, outer nuclear layer; INL, inner nuclear layer; GCL, ganglion cell layer; L, lens. Scale bar = 50 µm.

### RA regulates *her9* expression in the retina but Her9 is not required for RA’s effects on opsin expression

What regulates Her9 activity in the retina? Several previous studies have demonstrated that unlike many Hes/Hey/Her family members, Her9 does not respond to Notch signaling (Latimer et al., 2005; Leve et al., 2001; Reinhardt et al., 2015). However, it has been shown that Her9 is downstream of RA signaling in the developing inner ear (Radosevic et al., 2011). Given that RA signaling is critical for eye development (Hyatt et al., 1996a; Hyatt et al., 1996b) and has demonstrated effects on photoreceptor differentiation and opsin expression (Mitchell et al., 2015; Stenkamp et al., 2014; Stevens et al., 2011), we hypothesized that Her9 functions downstream of RA to regulate opsin expression and photoreceptor survival. We used in situ hybridization and qPCR on 36 hpf embryos to determine whether *her9* expression in the retina is altered by manipulation of the RA signaling pathway. Zebrafish embryos were treated in the dark from 24 to 36 hpf with 1 μM RA, 100 μM of the RA signaling inhibitor diethylaminobenzaldehyde (DEAB), or 0.3% DMSO alone as a carrier control. Embryos were then processed for WISH or qPCR to detect *her9* expression. In control embryos, *her9* expression was observed in the peripheral CMZ and around the lens and choroid fissure as described above (Fig. 12A). We observed a noticeable decrease in *her9* expression in the retina when embryos were treated with DEAB (Fig. 12A’). In contrast, there was a significant increase in the expression of *her9* all across the retina when embryos were exposed to RA (Fig. 12A”). qPCR analysis for *her9* expression confirmed the decrease and increase in *her9* expression following DEAB and RA treatments, respectively (Fig. 12B). From these data, we conclude that *her9* expression in the retina is regulated by RA signaling.

**Figure 12.**
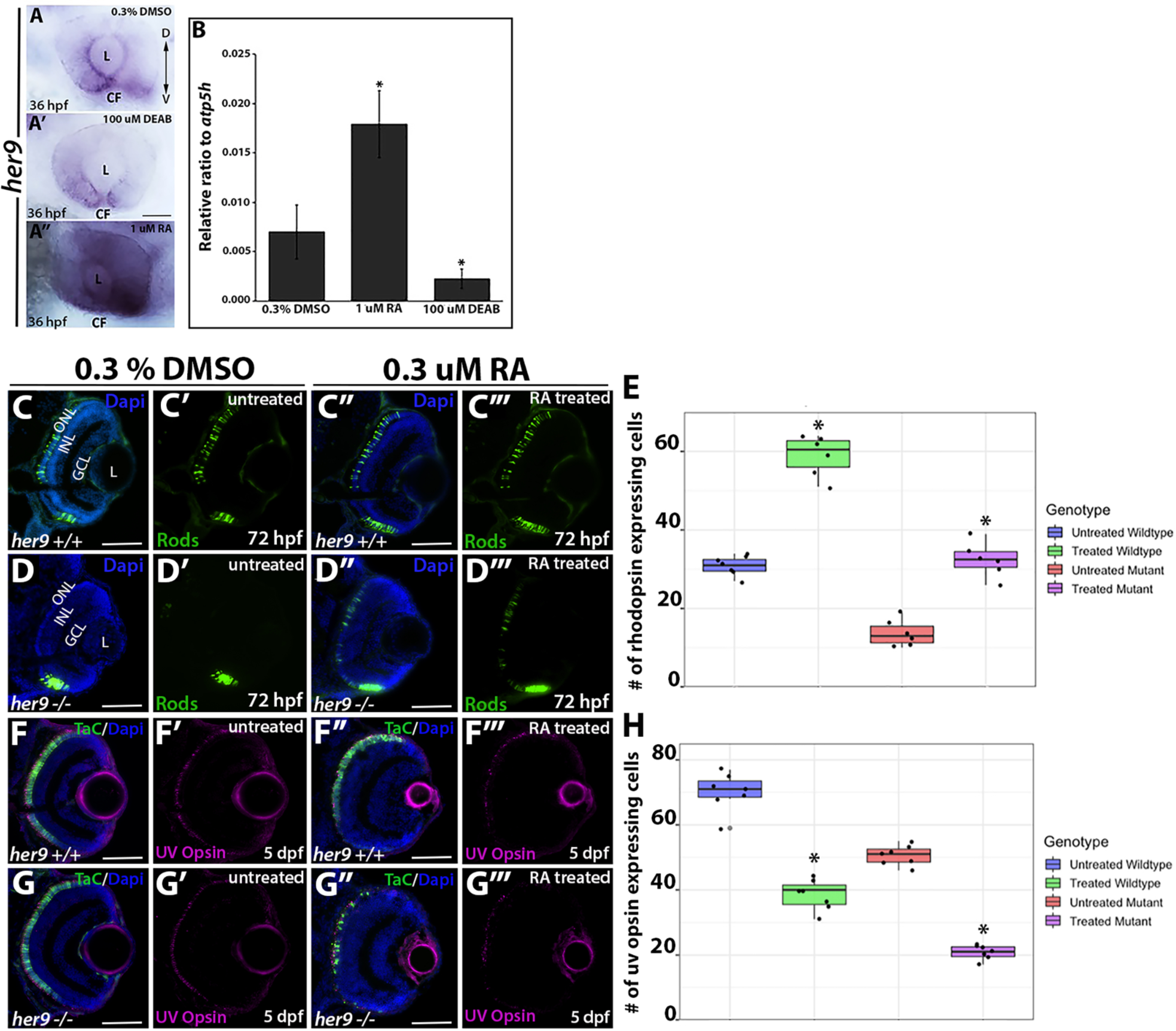
RA regulates *her9* expression but Her9 is not required for the effects of RA on opsin expression. WISH for *her9* expression at 36 hpf in control treated (**A**), DEAB treated (**A’**) and RA treated (**A”**) retinas. **(B)** qPCR for *her9* expression in heads of 36 hpf zebrafish embryos following drug treatments. N=20 heads per biological replicate, 3 biological replicates per drug treatment. *p<0.05. GFP expression in control treated vs. RA-treated XOPs:GFP;*her9+/+* (**C-C”’**) or XOPs:GFP;*her9-/-* (**D-D”’**) retinas. **(E)** Cell counts comparing rhodopsin expressing cells in the untreated and treated retinas. **(F-F”’)** IHC for UV opsin in control vs. RA-treated TαC:GFP;*her9+/+* (**F-F”’**) or TαC:GFP;*her9-/-* **(G-G”’)** retinas. **(H)** Cell counts comparing UV opsin expressing cells in the untreated and treated retinas. CF, choroid fissure; ONL, outer nuclear layer; INL, inner nuclear layer; GCL, ganglion cell layer; L, lens; Scale bar= 50µm

Previous work demonstrated that exposure to exogenous RA between 2 and 5 dpf resulted in increased rhodopsin expression and decreased red, UV, and blue cone opsin expression in zebrafish (Prabhudesai et al., 2005). To test the hypothesis that Her9 is required to mediate the effects of RA signaling on photoreceptor opsin expression, we treated WT and *her9* mutant zebrafish with RA from 24-48 hpf (for rods) or 24 hpf-5 dpf (for cones), and then examined XOPS:GFP or TαC:GFP transgene and cone opsin expression by IHC. In WT zebrafish, RA treatment resulted in a significant increase in GFP+ rhodopsin expressing cells dorsally and ventrally compared to the untreated retinas, and a significant decrease in UV and green cone opsin expression, as expected (Fig. 12 and Fig. S5). Surprisingly, an increase in GFP+ rhodopsin expressing cells and a decrease in UV and green cone opsin expressing cells were also observed in RA-treated *her9* mutants when compared to control treated mutants (Fig. 12 and Fig. S5). Taken together, these data suggest that although *her9* expression in the retina is downstream of RA signaling, Her9 is not strictly required for the effects of RA on rod and cone opsin expression.

## Discussion

The zebrafish *her9* gene is the ortholog of the human *HES4* gene and is one of the least studied bHLH-O transcription factors. This is largely due to the fact that a Hes4 homolog is absent from the mouse and rat genomes (El Yakoubi et al., 2012). Since *HES4* is expressed in humans and has been shown to play critical roles in the development of several vertebrate tissues, further investigations of Her9 function are needed. To our knowledge, ours is the first study to describe a germline null mutation of *her9*, which we believe will be a powerful resource for elucidating its role in several developmental contexts.

In this study, we focused on the role of Her9 during retinal development. We observed *her9* expression predominantly in the ventral most portion of the developing retina at 24 hpf, which localized to the ciliary marginal zone (CMZ) around 36 hpf until at least 5 dpf (Fig. 1). This expression pattern is consistent with what has previously been described for *her9* in Xenopus and Medaka (El Yakoubi et al., 2012; Reinhardt et al., 2015), suggesting a conserved mode of action in the developing retina at least across fish and amphibians.

To characterize the role of Her9 in zebrafish development, we used CRISPR/Cas to generate null mutations in *her9*. The mutations caused frame-shifts, and early termination codons, leading to nonsense-mediated decay of *her9* mRNA, which is supported by our sequencing, qPCR and Western blot data. The *her9* mutants appear developmentally delayed at around 20 hpf but by 72 hpf many *her9* mutant larvae look similar to their WT siblings. The relatively mild embryonic phenotype could be due to the buffering effects of maternally deposited WT *her9* mRNA. Nevertheless, by 5 dpf the mutants have not developed a swim bladder, display craniofacial and gastrointestinal defects, darker pigmentation, and abnormal swimming patterns. The *her9* mutants do not survive past 12 dpf, which may be due to problems with feeding and digestion.

One of the most striking phenotypes in the *her9* mutant consists of a significant decrease in rod and double cone photoreceptors, which is associated with elevated levels of apoptosis in the ONL and dysmorphic and truncated outer segments in the surviving rods and red/green cones. Intriguingly, this defect does not seem to affect the short wavelength (blue and UV) photoreceptor subtypes. The rod-cone dystrophy observed in *her9* mutants suggests that Her9 is required for rod and cone maintenance and survival. The degeneration of the rod photoreceptors seems more severe than the cones, supporting a rod-cone progression in degeneration. Given that we did not observe expression of *her9* in mature photoreceptors, this suggests that Her9 acts non-cell autonomously to maintain rods and double cones. It is unlikely that the RPE contributes to the photoreceptor phenotypes, since we did not observe *her9* expression in the RPE, nor were there any obvious RPE structural defects in *her9* mutants. One possibility is that one of the functions of Her9 in the peripheral retina is to influence gradients of retinoic acid or thyroid hormone signaling, which are known to be important for photoreceptor maintenance (see below). In any case, the truncation of the cone outer segments could explain the lack of VBA and OKR responses in the *her9* mutants, although we cannot rule out the possibility that the absent OKR response is due to defects in other tissues outside the retina. Given that we observed a significant decrease in rod and cone opsin expression, it is also possible that other phototransduction proteins are downregulated as well. Future electrophysiological and ultrastructural studies could help to resolve the functional consequences of Her9 loss to the photoreceptor outer segments.

The process of photoreceptor development has been extensively studied and important transcription factors that regulate photoreceptor specification have been identified, such as Crx, Nr2e3, and *thr*β, among many others. In our investigations of whether Her9 is required for rod and/or cone lineage specification, we saw no significant changes in *crx* expression in *her9* mutant retinas, indicating that Her9 is not required at this stage of photoreceptor specification. However, we did observe a significant decrease in expression of *Nr2e3* and *thr*β in *her9* mutants, indicating that Her9 may be required for the specification of some photoreceptor subtypes once progenitors have chosen a specific photoreceptor lineage.

The reduction in *thrβ* expression in *her9* mutants may contribute to additional phenotypes outside of the retina. Lui and Chan (Liu and Chan, 2002) showed that thyroid hormones are important for the embryonic to larval transition in zebrafish. Inhibition of Thrβ produced a missing swim bladder, defects in the gastrointestinal tract, and craniofacial defects. These phenotypes resemble those observed in the *her9* mutants where we also see a loss of swim bladder, gastrointestinal tract defects, and craniofacial abnormalities in addition to the photoreceptor defects. Furthermore, given that there is a direct link between *thrβ* expression and cone development and differentiation (Suzuki et al., 2013), our data suggest that Her9 could be directly or indirectly required for *thrβ* expression. This theory could be tested by exposing mutant embryos to L-thyroxine (T4) and 3,5,3’ –L-triiodothyronine (T3) to see whether we can rescue any components of the *her9* mutant phenotype.

In addition to decreases in photoreceptors, we also observed significant decreases in the number of Müller glia, ganglion cells, and in proliferating cells in the CMZ of *her9* mutants. Investigation into the role of Her9/Hes4 in both Medaka and *Xenopus* have demonstrated expression of *her9* in the CMZ of the developing and post-embryonic retina (El Yakoubi et al., 2012; Reinhardt et al., 2015). Consistent with those data, we also observed the expression of *her9* localized to the CMZ starting at 36 hpf. We observed a significant decrease in the proliferating cells of the CMZ of *her9* mutants that became more severe over time. This indicates that Her9 is required for the maintenance of proliferating retinal progenitor cells. Morpholino knockdown of Hes4 in Xenopus produced dose-dependent eye defects from small eyes to animals with no eyes at all (El Yakoubi et al., 2012). In our zebrafish *her9* mutants, we observed a variable degree of microphthalmia but never complete loss of eyes. This difference in phenotypic severity could be due to species-specific requirements for Her9 during oculogenesis, or to differences in experimental approach. In addition to a decrease in retinal cells, previous studies in *Xenopus* have shown that the loss of Hes4 leads to a significant increase in apoptosis in the retina (Nagatomo and Hashimoto, 2007; Nichane et al., 2008a; Nichane et al., 2008b). In our model, we also observe a significant increase in apoptosis in the retina and brain of our *her9* mutant which leads us to conclude that Her9 plays a role in cellular proliferation early on in retinal development and then cell survival later.

Radosevic et al. (2011) demonstrated that Her9 is downstream of the Retinoic acid (RA) signaling pathway in the neural patterning that underlies the development of the otic vesicle. With these data in mind, we wanted to determine whether Her9 is also downstream of RA signaling in the developing retina. Accordingly, we observed an increase in *her9* expression in the retina following exogenous treatment with RA, and blocking RA signaling with DEAB resulted in a decrease in *her9* expression. These data demonstrate that *her9* expression is regulated by RA signaling in the developing zebrafish retina. Taken together with the outer segment defects and decreased opsin expression in *her9* mutants, as well as previous studies demonstrating that opsin expression is regulated by RA signaling (Hyatt et al., 1996a; Hyatt et al., 1996b; Mitchell et al., 2015), we hypothesized that loss of Her9 was directly disrupting opsin expression downstream of RA signaling. However, when we exposed *her9* mutants to exogenous RA, we observed similar modulations in opsin expression to those observed on their WT siblings. This suggests that Her9 is not acting directly on opsin expression and may instead be acting downstream of RA to indirectly regulate photoreceptor differentiation and opsin expression. Future studies will include determining whether Her9 is interacting with other components of the RA pathway such as retinoic acid receptors, retinoid X receptors, or retinoic acid synthesis or degradation enzymes.

In summary, we have characterized a novel role for Her9 in photoreceptor development, maintenance, outer segment morphology and opsin expression. Given that the *her9* mutant displays such significant defects in rod and cone photoreceptors our results implicate Her9/HES4 as an important component of the gene regulatory network that regulates photoreceptor differentiation and survival. Future studies of Her9/HES4 could add to our understanding of its role in regulating retinal stem cell proliferation at the ciliary margin and determine how its loss leads to photoreceptor degeneration.

## Materials and methods

### Zebrafish lines and maintenance

All zebrafish lines were bred, housed, and maintained at 28.5°C on a 14 hour light: 10-hour dark cycle, in accordance with established protocols for zebrafish husbandry (Westerfield, 2000). The Tg(3.2TαC: EGFP) transgenic line (hereafter called TαC:EGFP) has been previously described (Kennedy et al., 2001), and was generously provided by Susan Brockerhoff (University of Washington, Seattle, WA). The Tg(gfap: GFP)mi2001 line (hereafter referred to as GFAP:GFP), was previously described (Bernardos and Raymond, 2006) and was obtained from the Zebrafish International Resource Center (ZIRC, Eugene, OR). The Tg(XlRho: EGFP) transgenic line (hereafter called XOPs:GFP) has been previously described (Fadool, 2003), and was obtained from James Fadool (Florida State University, Tallahassee, FL). The Tg(fli1a: EGFP)y1Tg transgenic line, (hereafter called fli1:GFP) has been previously described (Lawson and Weinstein, 2002), and was obtained from ZIRC. The Tg(atoh7:GFP) transgenic line, (hereafter called ath5:GFP) has been previously described (Masai et al., 2003) and was obtained from ZIRC. Embryos were anesthetized with Ethyl 3-aminobenzoate methanesulfonate salt (MS-222, Tricaine, Sigma-Aldrich, St. Louis, MO) and adults were euthanized by rapid cooling as previously described (Wilson et al., 2016). All animal procedures were carried out in accordance with guidelines established by the University of Kentucky Institutional Animal Care and Use Committee (IACUC) and the ARVO Statement for the Use of Animals in Ophthalmic and Vision Research.

### Generation of *her9* mutant zebrafish by CRISPR

The *her9* target sites and single strand DNA oligonucleotides used to generate the guide RNAs were selected using the ZiFit online tool (http://zifit.partners.org/ZiFiT/). The target sites for CRISPR/Cas9 genome editing were selected within the first and third exons of the *her9* gene. The first target site is 54 bp 3’ of the translation start site and 46 bp upstream of the beginning of the bHLH domain, while the second target site is immediately after the region corresponding to the Orange domain (sequences listed in Supplemental Table 2).

The pT3TS-nls-zCas9-nls (Addgene: 46757) expression vector was used to produce Cas9 mRNA. This vector contains a zebrafish codon-optimized Cas9 coding sequence flanked by the nuclear localization sequence. The plasmid was linearized using XbaI (NEB: R0145S) and purified by phenol:chloroform extraction and ethanol precipitation. Cas9 mRNA was generated using Ambion mMESSAGE mMACHINE T3 Transcription Kit (Life Technologies: AM1304) and purified by phenol:chloroform extraction and ethanol precipitation.

Screening for mutations in *her9* was performed by high resolution melting analysis (HRMA). Genomic DNA was collected from 24 hpf embryos as described below. HRMA was performed using the LightCycler 480 High-Resolution Melting Master (Roche) kit according to the manufacturer’s instructions on a LightCycler 96 Real-Time PCR System (Roche).

### Genomic DNA (gDNA) extraction and amplification

gDNA was extracted from whole embryos or from tail clips of adult fish. The embryos or tails were placed in 20 ul of 1x Thermopol buffer (NEB, Ipswich, MA) and incubated at 95° C to soften the tissue. The tissue was cooled on ice for 10 min before 5 ul of Proteinase K (Pro K; Sigma-Aldrich, St. Louis, MO) was added to each and incubated at 55° C overnight. After at least 18 hours of digestion, tubes containing the digested tissue were incubated at 95° C for 15 min to deactivate the Pro K. The gDNA was then amplified using primers described in Supplemental Table 1.

### Restriction fragment length polymorphism (RFLP) analysis

The mutant *her9* allele carrying a 1 bp deletion was identified by RFLP analysis due to the introduction of a BsaJI restriction site (NEB: R0563S). The mutant *her9* allele carrying a 1 bp insertion was identified by RFLP due to the introduction of a BfaI restriction site (NEB: R0568S). Genomic DNA from whole embryos or from tail clips was extracted and amplified by PCR as described above, then subjected to restriction digest. The restriction digests were visualized on a 2% agarose gel.

### mRNA synthesis and microinjections

The coding region of *her9* was amplified by RT-PCR and the cDNA was cloned into a pGEMT-easy plasmid (Promega, Madison, WI). Capped mRNA was synthesized using the mMessage T7 kit (Ambion, Austin, TX) according to the manufacturer’s instructions and purified by phenol-chloroform extraction and ethanol precipitation. The mRNA was injected at a volume of 4.18 nl/embryo (1.5 ng) in buffered solution with 0.025% Dextran red into the yolk of 1-cell stage zebrafish.

### Western Blot Analysis

Protein lysate was extracted from 20 heads of 3 dpf *her9* wildtype and mutant larvae. Protein was quantified by Bradford Assay (BIO-Rad), and 30 μg was diluted 1:3 in gel loading buffer (NEB), sonicated, spun down, then separated by SDS-PAGE on Mini-PROTEAN TGX Precast Gels (BIO-Rad). Following gel electrophoresis, separated proteins were transferred to a nitrocellulose membrane. The membrane was blocked with 5% non-fat milk/PBST for 30 minutes at room temperature then incubated in anti-HES4 (rabbit polyclonal, 1:500, Thermo Fisher) or anti-β actin (mouse monoclonal, 1:1000, Santa Cruz)/PBST solution overnight at 4°C. Membranes were washed in PBST then incubated in secondary antibody for one hour at room temperature, developed with HRP Development Reagent (BIO-Rad) for 1 minute, and imaged with a Chemidoc Bioimager (BIO-Rad).

### RNA isolation and riboprobe synthesis

RNA was isolated from whole embryos at indicated developmental time points. RNA was extracted from pools of embryos using TRIzol Reagent (Life Technologies, Invitrogen), following the manufacturer’s protocol, then treated with RNAse-Free DNAse I (Roche, Indianapolis, IN) to remove genomic DNA as previously described (Wilson et al., 2009). For riboprobes, PCR products from the unique regions of *her9* were cloned into the pGEMT-easy vector (Promega, Madison, WI). Plasmids were linearized using either NcoI and SpeI restriction enzymes (NEB, Ipswich, MA). Riboprobes were synthesized from the plasmids by in vitro transcription using either T7 or Sp6 polymerase and a digoxigenin (DIG) labeling kit (Roche Applied Science, Indianapolis, IN). The primer sequences used for riboprobe preparation are given in Supplemental Table 1.

### RT-PCR and real-time quantitative RT-PCR (qPCR)

The GoScript Reverse Transcriptase System (Promega, Madison, WI) was used to synthesize the first strand cDNA from 1μg of the extracted RNA. PCR primers were designed to amplify unique regions of the *her9* and *atp5h* cDNAs (Eurofins Genomics; www.eurofinsgenomics.com; Supplemental Table 1). Faststart Essential DNA Green Master mix (Roche) was used to perform qPCR on a Lightcycler 96 Real-Time PCR System (Roche). The relative transcript abundance was normalized to *atp5h* expression as the housekeeping gene control (Wen et al., 2015), and was calculated as fold-change relative to 36 hours post fertilization (hpf) for developmental expression, and fold-change relative to wild type siblings, untreated, or uninjected embryos for the mRNA rescue and RA experiments. RT-PCR and qPCR experiments were performed with three biological replicates and three technical replicates. RT-PCR was performed on a Mastercycler Pro thermocycler (Eppendorf, Westbury, NY). PCR products were visualized on a 1% agarose gel.

### Visually mediated background adaptation assay (VBA assay)

The VBA assay was adapted from previously described protocols (Mueller and Neuhauss, 2014; Viringipurampeer et al., 2014). Embryos at 5 dpf were incubated in the dark for 2 hours and the dorsal cranial pigment was immediately imaged after removal from the dark. The embryos were then placed under ambient light for 30 minutes. The dorsal cranial pigment was imaged immediately following light exposure. Images of the heads of the embryos were scored by naïve observers as either “Light” or “Dark”. All imaged embryos were then genotyped by RFLP analysis. The assay was performed a minimum of three times. In each assay, 10 embryos were screened. Results were analyzed by Chi-square test.

### Optokinetic response assay (OKR assay)

Embryos were collected at 5 dpf and mounted in a 35-mm petri dish filled with 5% methylcellulose. The embryos were mounted near the surface of the methylcellulose, dorsal side up. The petri dish was then placed inside of a rotating drum containing illuminated vertical stripes. The drum was rotated clockwise for 30 seconds (s) at 8 rpm then 30 s counterclockwise at 8 rpm and finally clockwise for another 30 s. Embryos were scored “responders” if they displayed smooth saccade eye movements in response to the drum rotation in the direction of rotation for at least two of the 30 second intervals. Embryos were scored as “non-responders” if they displayed no eye movement in any of the 30 s intervals. After scoring, the embryos were collected in 1x Thermopol buffer for gDNA extraction and genotyping using RFLP analysis as described above. The OKR assay was performed a minimum of three times, and at least 10 embryos were scored each time.

### Mobility Assay

Single 5 dpf zebrafish larvae were filmed for 1 minute in a 96-well plate to assess their swimming behavior. Each larva was subsequently screened for the *her9* 1bp insertion via PCR amplification and a restriction digest with BfaI (NEB: R0568S). The videos were linked to the appropriate genotype and analyzed using EthoVision software (www.noldus.com/animal-behavior-research/products/ethovision-xt). EthoVision analysis produced individual tracks for each larva, a compiled heat map of movement by genotype, and comparisons of total velocity and distance traveled.

### Whole-mount and fluorescent in situ hybridization (WISH and FISH)

Embryos were manually dechorionated, collected at selected developmental time points (18, 24, 48, 72, and 120 hpf) and fixed as described above. WISH was performed as previously described (Forbes-Osborne et al., 2013). Solutions were prepared with diethyl pyrocarbonate (DEPC)-treated water. DIG-labeled riboprobes (3 ng/μl) were hybridized to the samples overnight at 60°C in hybridization buffer. After washing and blocking, samples were incubated overnight at 4°C with an anti-DIG-AP antibody (Roche, diluted 1:2000 in blocking solution). The next day, the embryos were washed and equilibrated in NTMT buffer followed by coloration with 4-nitro blue tetrazolium (NBT, Roche) and 5-Bromo-4chloro-3-indolyl-phosphate, 4-toluidine salt (BCIP, Roche) in NTMT buffer. A stop solution (PBS pH 5.5, 1mM EDTA) was used to end the coloration reaction and embryos were placed in 40% glycerol for imaging.

FISH was performed as previously described (Pillai-Kastoori et al., 2014). Embryos were fixed and sectioned as previously described (Coomer and Morris, 2018). Sections were post-fixed in 1% PFA and rehydrated in PBST. The TSA plus Cy3 Kit (Perkin Elmer Inc, Waltham, MA) was used for probe detection. The sections were counterstained with 4’, 6-diamidino-2-phenylindole (DAPI, 1:10,000 dilution, Sigma-Aldrich), mounted in 60% glycerol, and imaged on an inverted fluorescent microscope (Nikon Eclipse Ti-U, Nikon Instruments, Melville, NY) using a 20x or 40x objective and a Leica SP8 DLS confocal/digital light sheet system (Leica Biosystems, Nussloch, Germany) using a 40x or 63x objective. At least 10 embryos and 10 sections were analyzed for each time point and probe.

### Immunohistochemistry

Immunohistochemistry was performed as previously described (Coomer and Morris, 2018). The following primary antibodies were used: 4C12 (mouse, 1:100) generously provided by J. Fadool (Florida State University, Tallahassee, FL), which labels rod photoreceptor cell bodies; 1D1 (mouse, 1:100, J. Fadool, FSU, Tallahassee, FL), which recognizes rhodopsin; Zpr-1 (mouse, 1:20, ZIRC), which labels red-green double cones; HuC/D (mouse, 1:20, Invitrogen, Grand Island, NY), which recognizes retinal ganglion cells and amacrine cells; Prox1 (rabbit, 1:2000, Millipore, Billerica, MA), which recognizes horizontal cells; anti-PCNA (mouse, 1:100, Santa Cruz Biotechnology, Dallas, TX), which labels proliferating cells; Zpr-3 (rabbit, 1:1000, ZIRC), which labels double cone and rod photoreceptor outer segments; and anti-Blue, -green, and -UV Opsin (rabbit, 1:1,000), generously provided by D. Hyde (University of Notre Dame) which labels the blue, green, and UV cones, respectively. Alexa Fluor 488 goat anti-mouse, 488 goat anti-rabbit, 546 goat anti-rabbit, and 546 goat anti-mouse secondary antibodies (Molecular Probes, Invitrogen) were all used at 1:200 dilution. Nuclei were visualized by counterstaining with DAPI (1:10,000 dilution). Samples were mounted in 60% glycerol in PBS. Images were taken at 20x and 40x on an inverted fluorescent microscope and a Leica SP8 DLS confocal/digital light sheet system using a 40x or 63x objective. Ten sections were analyzed on each slide and for each antibody.

### TUNEL staining

Terminal deoxynucleotide transferase (TdT)-mediated dUTP nick end labeling (TUNEL) was performed on frozen retinal cryosections using the ApopTag Fluorescein Direct In Situ Apoptosis Detection Kit (Millipore, Billerica, MA). TUNEL staining was performed according to the manufacturer’s protocol. Images were taken at 20x and 40x on an inverted fluorescent microscope (Eclipse Ti-U, Nikon instruments). Ten sections were analyzed on each slide.

### RA treatment

Stocks of all-trans-RA in dimethyl-sulfoxide (DMSO) were thawed before use and diluted to 1 µM retinoic acid with 0.3% DMSO in embryo medium, 100 µM of the RA signaling inhibitor diethylaminobenzaldehyde (DEAB) in 0.3% DMSO in embryo medium or 0.3% DMSO alone as the carrier control. Fli1:EGFP embryos and embryos from *her9* heterozygous incrosses were immersed in the RA solution starting at 24 hpf and were incubated for 12 hours in complete darkness. At 36 hpf the embryos were rinsed in artificial fish water for 10 min and processed for immunohistochemistry, or the heads were removed and processed for qPCR.

To observe effects of RA on rod and cone photoreceptors, embryos from wild type or *her9* heterozygous incrosses were incubated in 0.3 μM RA solution starting at 48 hpf and the solution was refreshed every 24 hpf. The embryos were collected at 72 hpf for analysis of rods, and at 120 hpf for analysis of cones. The heads were removed and sectioned for IHC and imaging.

### Statistical analysis

Statistical analysis was performed using a Student’s t-test or Chi-square test with GraphPad Prism software (www.graphpad.com). For comparing the number of retinal cell types and other phenotypic features 10 wild type and 10 mutant animals were examined. All graph data are represented as the mean ± the standard deviation (s.d.).

## Acknowledgments

The authors would like to thank Lucas Vieira Francisco for excellent zebrafish care. We also thank Dr. David Hyde (University of Notre Dame) for generously providing the zebrafish opsin antibodies.

## References

Bae, Y. K., Shimizu, T. and Hibi, M. (2005). Patterning of proneuronal and inter-proneuronal domains by hairy- and enhancer of split-related genes in zebraf ish neuroectoderm. Development 132, 1375–1385.

Bernardos, R. L. and Raymond, P. A. (2006). GFAP transgenic zebrafish. Gene expression patterns : GEP 6, 1007–1013.

Brockerhoff, S. E. (2006). Measuring the optokinetic response of zebrafish larvae. Nature Protocols 1, 2448–2451.

Chen, J. C., Rattner, A. and Nathans, J. (2005). The rod photoreceptor-specific nuclear receptor Nr2e3 represses transcription of multiple cone-specific genes. Journal of Neuroscience 25, 118–129.

Coomer, C. E. and Morris, A. C. (2018). Capn5 Expression in the Healthy and Regenerating Zebrafish Retina. Investigative Ophthalmology & Visual Science 59, 3643–3654.

Delidakis, C., Monastirioti, M. and Magadi, S. S. (2014). E(spl): Genetic, Developmental, and Evolutionary Aspects of a Group of Invertebrate Hes Proteins with Close Ties to Notch Signaling. In Bhlh Transcription Factors in Development and Disease (ed. R. Taneja), pp. 217–262. San Diego: Elsevier Academic Press Inc.

Doerre, G. and Malicki, J. (2002). Genetic analysis of photoreceptor cell development in the zebrafish retina. Mechanisms of Development 110, 125–138.

El Yakoubi, W., Borday, C., Hamdache, J., Parain, K., Tran, H. T., Vleminckx, K., Perron, M. and Locker, M. (2012). Hes4 Controls Proliferative Properties of Neural Stem Cells During Retinal Ontogenesis. Stem Cells 30, 2784–2795.

Fadool, J. M. (2003). Development of a rod photoreceptor mosaic revealed in transgenic zebrafish. Developmental Biology 258, 277–290.

Farre, A., Mackin, R. and Stenkamp, D. L. (2019). Thyroid hormone regulates the tandemly-quadruplicated rh2 cone opsin gene array in zebrafish. Investigative Ophthalmology & Visual Science 60, 2.

Forbes-Osborne, M. A., Wilson, S. G. and Morris, A. C. (2013). Insulinoma-associated 1a (Insm1a) is required for photoreceptor differentiation in the zebrafish retina. Developmental Biology 380, 157–171.

Holzschuh, J., Wada, N., Wada, C., Schaffer, A., Javidan, Y., Tallafuss, A., Bally-Cuif, L. and Schillili, T. F. (2005a). Requirements for endoderm and BMP signaling in sensory neurogenesis in zebrafish. Development 132, 3731–3742.

Holzschuh, J., Wada, N., Wada, C., Schaffer, A., Javidan, Y., Tallafuss, A., Bally-Cuif, L. and Schilling, T. F. (2005b). Requirements for endoderm and BMP signaling in sensory neurogenesis in zebrafish. Development (Cambridge, England) 132, 3731–3742.

Hyatt, G. A., Schmitt, E. A., Fadool, J. M. and Dowling, J. E. (1996a). Retinoic acid alters photoreceptor development in vivo. Proceedings of the National Academy of Sciences of the United States of America 93, 13298–13303.

Hyatt, G. A., Schmitt, E. A., MarshArmstrong, N., McCaffery, P., Drager, U. C. and Dowling, J. E. (1996b). Retinoic acid establishes ventral retinal characteristics. Development 122, 195–204.

Il, L. L. N., Alur, R. P., Boobalan, E., Sergeev, Y. V., Caruso, R. C., Stone, E. M., Swaroop, A., Johnson, M. A. and Brooks, B. P. (2010). Two Novel CRX Mutant Proteins Causing Autosomal Dominant Leber Congenital Amaurosis Interact Differently With NRL. Human Mutation 31, E1472–E1483.

Kennedy, B. N., Vihtelic, T. S., Checkley, L., Vaughan, K. T. and Hyde, D. R. (2001). Isolation of a zebrafish rod opsin promoter to generate a transgenic zebrafish line expressing enhanced green fluorescent protein in rod photoreceptors. Journal of Biological Chemistry 276, 14037–14043.

Latimer, A. J., Shin, J. and Appel, B. (2005). her9 promotes floor plate development in zebrafish. Developmental Dynamics 232, 1098–1104.

Lawson, N. D. and Weinstein, B. M. (2002). In vivo imaging of embryonic vascular development using transgenic zebrafish. Developmental biology 248, 307–318.

Leve, C., Gajewski, M., Rohr, K. B. and Tautz, D. (2001). Homologues of c-hairy1 (her9) and lunatic fringe in zebrafish are expressed in the developing central nervous system, but not in the presomitic mesoderm. Development Genes and Evolution 211, 493–500.

Li, J., Zhang, B. B., Ren, Y. G., Gu, S. Y., Xiang, Y. H., Huang, C. and Du, J. L. (2015). Intron targeting-mediated and endogenous gene integrity-maintaining knockin in zebrafish using the CRISPR/Cas9 system. Cell Research 25, 634–637.

Liu, Y. W. and Chan, W. K. (2002). Thyroid hormones are important for embryonic to larval transitory phase in zebrafish. Differentiation 70, 36–45.

Liu, Z. H., Dai, X. M. and Du, B. (2015). Hes1: a key role in stemness, metastasis and multidrug resistance. Cancer Biology & Therapy 16, 353–359.

Livesey, F. J. and Cepko, C. L. (2001). Vertebrate neural cell-fate determination: Lessons from the retina. Nature Reviews Neuroscience 2, 109–118.

Masai, I., Lele, Z., Yamaguchi, M., Komori, A., Nakata, A., Nishiwaki, Y., Wada, H., Tanaka, H., Nojima, Y., Hammerschmidt, M., et al. (2003). N-cadherin mediates retinal lamination, maintenance of forebrain compartments and patterning of retinal neurites. Development 130, 2479–2494.

Mitchell, D. M., Stevens, C. B., Frey, R. A., Hunter, S. S., Ashino, R., Kawamura, S. and Stenkamp, D. L. (2015). Retinoic Acid Signaling Regulates Differential Expression of the Tandemly-Duplicated Long Wavelength-Sensitive Cone Opsin Genes in Zebrafish. Plos Genetics 11, 33.

Morris, A. C., Forbes-Osborne, M. A., Pillai, L. S. and Fadool, J. M. (2011). Microarray Analysis of XOPS-mCFP Zebrafish Retina Identifies Genes Associated with Rod Photoreceptor Degeneration and Regeneration. Investigative Ophthalmology & Visual Science 52, 2255–2266.

Morrow, D., Cullen, J. P., Liu, W., Guha, S., Sweeney, C., Birney, Y. A., Collins, N., Walls, D., Redmond, E. M. and Cahill, P. A. (2009). Sonic Hedgehog induces Notch target gene expression in vascular smooth muscle cells via VEGF-A. Arteriosclerosis, thrombosis, and vascular biology 29, 1112–1118.

Mueller, K. P. and Neuhauss, S. C. F. (2014). Sunscreen for Fish: Co-Option of UV Light Protection for Camouflage. Plos One 9, 5.

Muller, M., vonWeizsacker, E. and CamposOrtega, J. A. (1996). Expression domains of a zebrafish homologue of the Drosophila pair-rule gene hairy correspond to primordia of alternating somites. Development 122, 2071–2078.

Nagatomo, K. I. and Hashimoto, C. (2007). Xenopus hairy2 functions in neural crest formation by maintaining cells in a mitotic and undifferentiated state. Developmental Dynamics 236, 1475–1483.

Neuhauss, S. C. F., Biehlmaier, O., Seeliger, M. W., Das, T., Kohler, K., Harris, W. A. and Baier, H. (1999). Genetic disorders of vision revealed by a behavioral screen of 400 essential loci in zebrafish. Journal of Neuroscience 19, 8603–8615.

Nichane, M., de Croze, N., Ren, X., Souopgui, J., Monsoro-Burq, A. H. and Bellefroid, E. J. (2008a). Hairy2-Id3 interactions play an essential role in Xenopus neural crest progenitor specification. Developmental Biology 322, 355–367.

Nichane, M., Ren, X., Souopgui, J. and Bellefroid, E. J. (2008b). Hairy2 functions through both DNA-binding and non DNA-binding mechanisms at the neural plate border in Xenopus. Developmental Biology 322, 368–380.

Pillai-Kastoori, L., Wen, W., Wilson, S. G., Strachan, E., Lo-Castro, A., Fichera, M., Musumeci, S. A., Lehmann, O. J. and Morris, A. C. (2014). Sox11 Is Required to Maintain Proper Levels of Hedgehog Signaling during Vertebrate Ocular Morphogenesis. Plos Genetics 10, 19.

Prabhudesai, S. N., Cameron, D. A. and Stenkamp, D. L. (2005). Targeted effects of retinoic acid signaling upon photoreceptor development in zebrafish. Developmental Biology 287, 157–167.

Radosevic, M., Robert-Moreno, À., Coolen, M., Bally-Cuif, L. and Alsina, B. (2011). Her9 represses neurogenic fate downstream of Tbx1 and retinoic acid signaling in the inner ear. Development (Cambridge, England) 138, 397–408.

Rath, M. F., Morin, F., Shi, Q., Klein, D. C. and Moller, M. (2007). Ontogenetic expression of the Otx2 and Crx homeobox genes in the retina of the rat. Experimental Eye Research 85, 65–73.

Reinhardt, R., Centanin, L., Tavhelidse, T., Inoue, D., Wittbrodt, B., Concordet, J. P., Martinez-Morales, J. R. and Wittbrodt, J. (2015). Sox2, Tlx, Gli3, and Her9 converge on Rx2 to define retinal stem cells in vivo. Embo Journal 34, 1572–1588.

Stenkamp, D. L., Stevens, C. B., Frey, R. A. and Kawamura, S. (2014). Retinoic acid signaling regulates expression of the tandemly duplicated LWS1 and LWS2 genes in zebrafish. Investigative Ophthalmology & Visual Science 55, 3.

Stevens, C. B., Cameron, D. A. and Stenkamp, D. L. (2011). Plasticity of photoreceptor-generating retinal progenitors revealed by prolonged retinoic acid exposure. Bmc Developmental Biology 11, 25.

Suzuki, S. C., Bleckert, A., Williams, P. R., Takechi, M., Kawamura, S. and Wong, R. O. L. (2013). Cone photoreceptor types in zebrafish are generated by symmetric terminal divisions of dedicated precursors. Proceedings of the National Academy of Sciences of the United States of America 110, 15109–15114.

Takke, C., Dornseifer, P., Von Weizsacker, E. and Campos-Ortega, J. A. (1999). her4, a zebrafish homologue of the Drosophila neurogenic gene E(spl), is a target of NOTCH signalling. Development 126, 1811–1821.

Viringipurampeer, I. A., Shan, X., Gregory-Evans, K., Zhang, J. P., Mohammadi, Z. and Gregory-Evans, C. Y. (2014). Rip3 knockdown rescues photoreceptor cell death in blind pde6c zebrafish. Cell Death and Differentiation 21, 665–675.

Wen, W., Pillai-Kastoori, L., Wilson, S. G. and Morris, A. C. (2015). Sox4 regulates choroid fissure closure by limiting Hedgehog signaling during ocular morphogenesis. Developmental Biology 399, 139–153.

Wilson, J. M., Bunte, R. M. and Carty, A. J. (2009). Evaluation of Rapid Cooling and Tricaine Methanesulfonate (MS222) as Methods of Euthanasia in Zebrafish (Danio rerio). Journal of the American Association for Laboratory Animal Science 48, 785–789.

Wilson, S. G., Wen, W., Pillai-Kastoori, L. and Morris, A. C. (2016). Tracking the fate of her4 expressing cells in the regenerating retina using her4:Kaede zebrafish. Experimental Eye Research 145, 75–87.

