## Supplemental information for "Her9/Hes4 is required for retinal photoreceptor development, maintenance, and survival"

**Supplementary Information**

**Supplemental Figures and Legends**

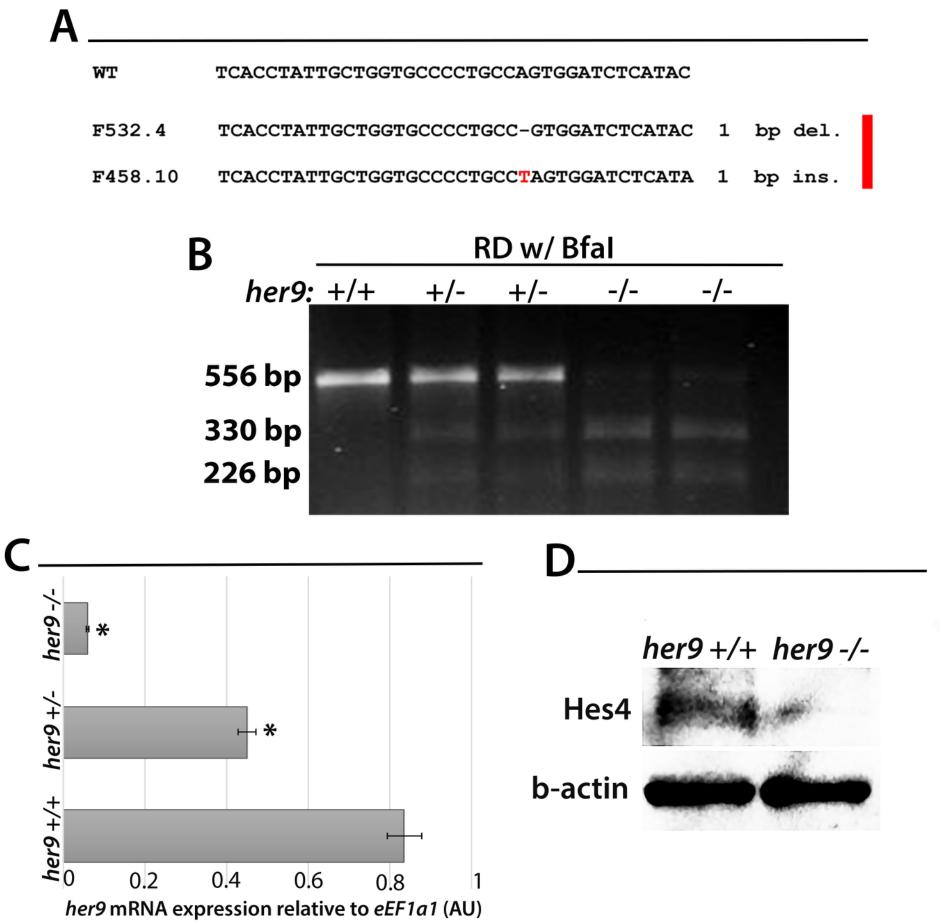


**Figure S1. Generation of *her9* mutants using CRISPR/Cas9. (A)** Comparison of WT *her9* sequence with 1bp deletion and 1bp insertion mutations. (**B)** RFLP analysis of WT, heterozygous and homozygous *her9* 1 bp insertion mutant cut with BfaI. (**C)** qPCR analysis of *her9* mRNA expression in WT, heterozygous and homozygous mutants at 48 hpf. (**D)** Western blot with a HES4 antibody, indicating the loss of Her9 protein in the *her9* mutants compared to WT siblings.

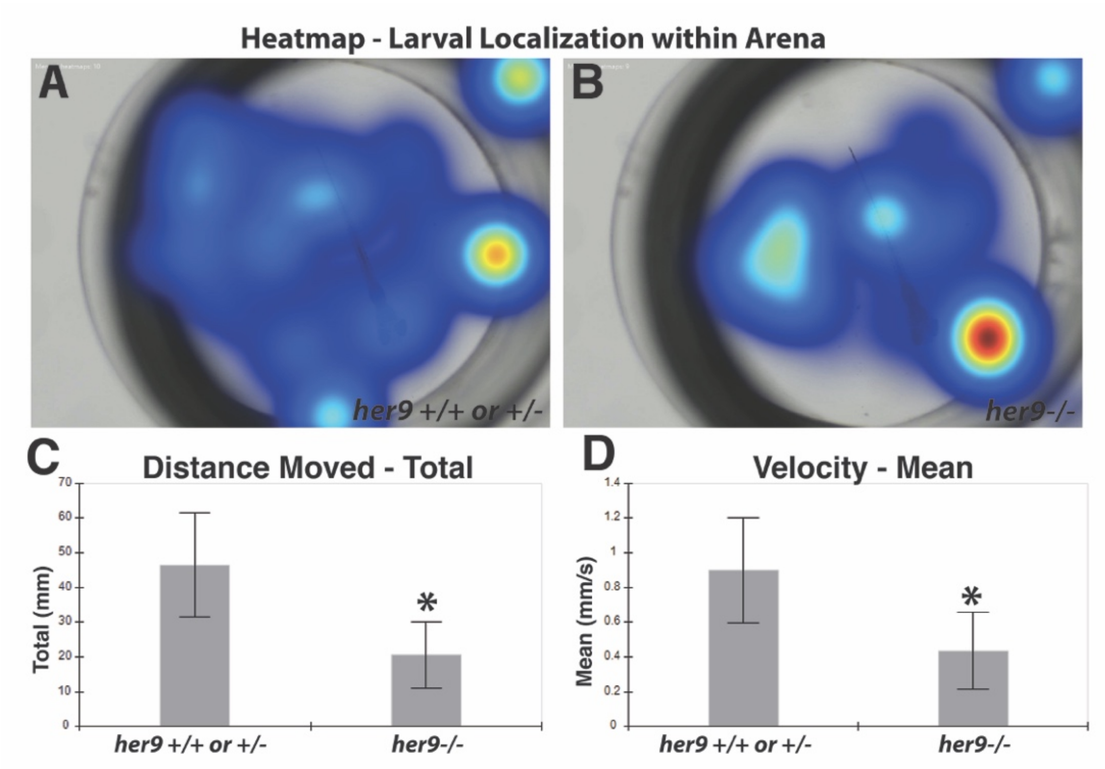


**Figure S2. Mobility assay of 5 dpf larvae. (A)** Heat map indicating the amount of time the WT or *her9*+/- larva spends in different parts of the arena. **(B)** Heat map indicating the amount of time the *her9*-/- larva spends in different parts of the arena. **(C)** Comparison of the average total distance travel by the larvae. The WT or *her9* +/- average total distance traveled was 46.47±14.91 mm, the *her9* -/- average total distance traveled was 20.59±9.59 mm. **(D)** Comparison of the average velocity of larvae. The WT or *her9*+/- average velocity was 0.901±0.302mm/s, the *her9*-/- average velocity was 0.436±0.22 mm/s.

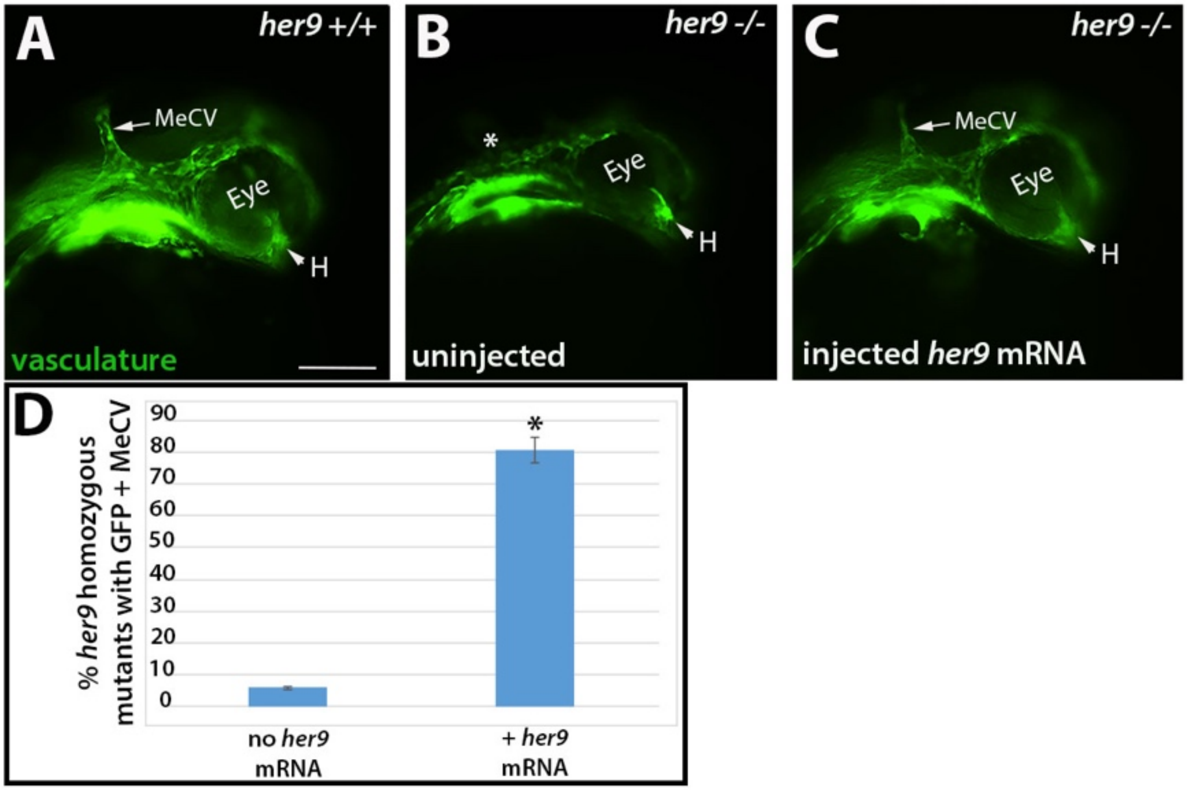


**Figure S3. Injection of *her9* mRNA rescues the *her9* mutant phenotype. (A)** Fli1:GFP+ midcerebral vein in WT embryos at 24 hpf. (**B)** Missing midcerebral vein in uninjected *her9* homozygous mutant embryos at 24 hpf (asterisk). (**C)** In *her9* homozygous mutants injected with *her9* mRNA, the midcerebral vein is now visible at 24 hpf. (**D)** Quantification of rescue of *her9* mutant phenotype after *her9* mRNA injections. MeCV, Midcerebral vein; H, hyaloid vein. Scale bar= 50 μm.

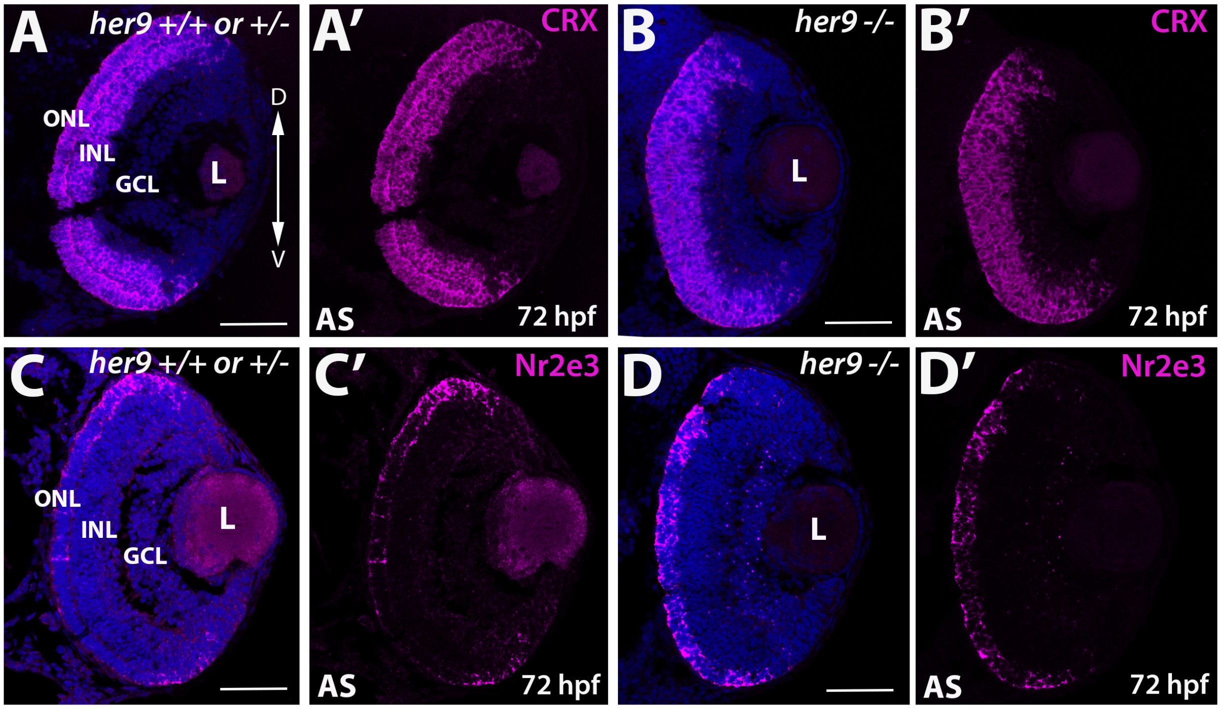


**Figure S4. Expression of *crx* and *Nr2e3* at 72 hpf.** (**A-B)** Fluorescent in situ hybridization (FISH). *Crx* expression at 72 hpf in WT and *her9* mutant retina. *Her9* mutants displayed similar expression to WT. **(C-D')** *Nr2e3* expression at 72 hpf in WT and *her9* mutant retina. Expression in the mutant ONL was distorted compared to the WT. L, lens; ON, optic nerve. Scale bar= 50 µm

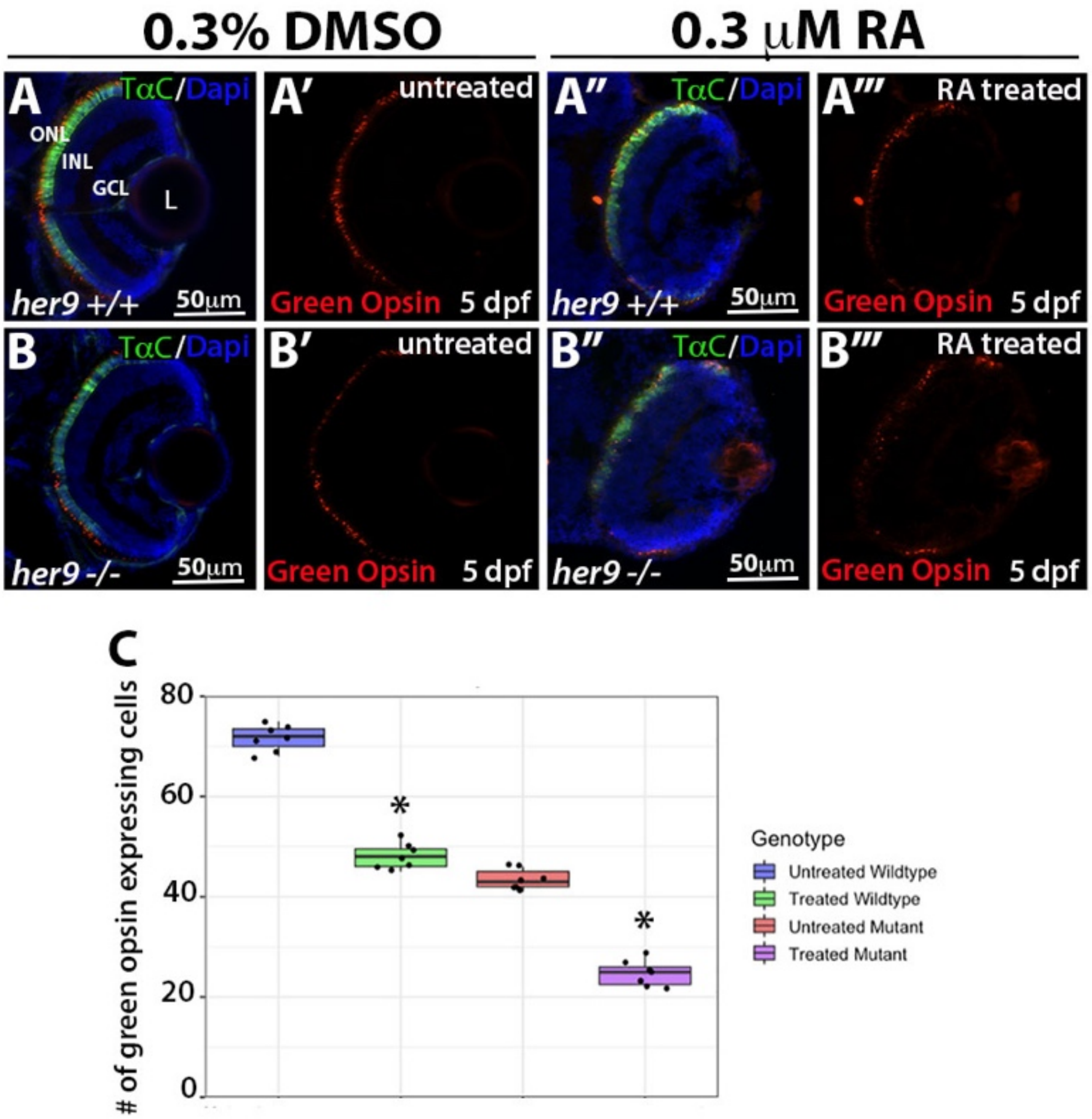


**Figure S5. Her9 is not required for the effects of RA on opsin expression. (A-A''')** IHC on TαC:GFP WT retinas comparing Green opsin expression after being treated with RA from 24- 120 hpf. **(B-B''')** IHC on TαC:GFP mutant retinas comparing Green opsin expression after being treated with RA from 24- 120 hpf. **(C)** Cell counts comparing Green opsin expressing cells in the untreated and treated retinas. ONL, outer nuclear layer; INL, inner nuclear layer; GCL, ganglion cell layer; L, lens; Scale bar= 50μm

| Primers | Sequences | Use |
|---|---|---|
| *her9* F | CCTGACGGAGAACTGAACACAAGACACACA | RT-PCR/ISH |
| *her9* R | TTTCTCAATGGTACGGCGGGTGCTCTGGGC | RT-PCR/ISH |
| *atp5h* F | TGCCATCTCAGCAAAACTTG | RT-PCR/qPCR |
| *atp5h* R | CACAGGCTCAGGAACAGTCA | RT-PCR/qPCR |
| *her9 CR* F | AAGCTTCCTGACGGAGAACTGAACACAAGACACACA | cDNA plasmid |
| *her9 CR* R | GAATTCTTTCTCAATGGTACGGCGGGTGCTCTGGGC | cDNA plasmid |
| *her9* F | CTCAGCAAGTACCGCGCAGGAT | qPCR |
| *her9* R | TCCCATACAACCGGACAGGTGG | qPCR |
| *tr*β*2* F | ACAGGGAGACTGTAGAGGTCTGA | ISH |
| *tr*β*2* R | TCAGTCTTCAAACACTTCCAGGAAG | ISH |
| *tr*β*2* F | AAGAGGGGCTCTGGCTCTTA | qPCR |
| *tr*β*2* R | GTTGTCCACAGACTCGCTGA | qPCR |
| *Nr2e3* F | CCAGCAGTGGGAAACACTAT | qPCR |
| *Nr2e3* R | ATGGGCTTTATCCACAGGAC | qPCR |
| *Nr2e3* F | ATCATCGCGGCCGCCACCATGGAGGATCCGATGTCAGAAATG | ISH |
| *Nr2e3* R | ACTACTAAGCTTATCAGTTTTGAACATGTCACACAAG | ISH |
| *crx* F | ATGCTGTGAACGGGTTAAC | qPCR |
| *crx* R | AAGCTTCCAGAATGTCCAG | qPCR |
| *rho* F | GCCTATGTCCAATGCCACCGG | qPCR |
| *rho* R | AGTTGACGGGGAAGCCGGTGA | qPCR |
| *LWS* F | GGCGAGGGGACGAAACAACA | qPCR |
| *LWS* R | CCATCGAGGGGCAATGTGGT | qPCR |
| *RH2* F | TCTACATCCCCATGTCAAACAGGA | qPCR |
| *RH2* R | AGTCAGGACATTGATGGGAAATCC | qPCR |
| *SWS1* F | CGTTCAATTCGGAAATGCTTCC | qPCR |
| *SWS1* R | TGTGCCCACGATAAATACAAAGCC | qPCR |
| *SWS2* F | AGCAAACGCCAGAACTGTTCGA | qPCR |
| *SWS2* R | CATGAATACGCCACTGTGTCCCA | qPCR |
| *eEF1*α F | CTTCTCAGGCTGACTGTGC | qPCR |
| *eEF1*α R | CCGCTAGCATTACCCTCC | qPCR |
| *six7* F | ACCGGTTTACACGCGAATCT | qPCR |
| *six7* R | GCAGGGGGAACTTCTTACGG | qPCR |
| *gdf6a* F | TATTGGTGGTCTACACGCGG | qPCR |
| *gdf6a* R | TAAGCGCAGTTCTCCGTCTG | qPCR |
| *vsx2* F | GAAGTGTTGGGGTCGCAGTA | qPCR |
| *vsx2* R | GTGCATCCCTAGAAGCCAGG | qPCR |

**Supplemental Table 1.** Primer sequences used for RT-PCR and qPCR. RT-PCR primers were also used to design WISH and FISH probes. F, forward; R, reverse; CR, coding region.

| Oligo name | Oligo sequence | Use |
|---|---|---|
| *her9* F CRISPR1 | GTGTTTGATCATGCCAGCCGAT | HRMA/sequencing |
| *her9* R CRISPR1 | CCTTTCTATGCTCGCTGGCATT | HRMA/sequencing |
| *her9* F CRISPR2 | CCACCTGTCCGGTTGTATGGG | HRMA/sequencing |
| *her9* R CRISPR2 | GAGGCGCCGTTGATGGGTAAC | HRMA/sequencing |
| *her9* C1 F | ATCTATAAATACCGCTGGCGTGTGG | RFLP |
| *her9* C1 R | TCTCGTTGATTCTCGCTCTGCG | RFLP |
| CRISPR1 target site | GGAGTATGAGATCCACTGGC | *her9* target 1 |
| CRISPR1 oligo-1 | TAGGAGTATGAGATCCACTGGC | *her9 C1 oligo-1* |
| CRISPR1 oligo-2 | AAACGCCAGTGGATCTCATACT | *her9 C1 oligo-2* |
| CRISPR2 target site | GGCAGGCTGAGGGTAGTTCA | *her9 target 2* |
| CRISPR2 oligo-1 | TAGGCAGGCTGAGGGTAGTTCA | *her9 C2 oligo-1* |
| CRISPR2 oligo-2 | AAACTGAACTACCCTCAGCCTG | *her9 C2 oligo-2* |

**Supplemental Table 2.** Oligo sequences used for her9 CRIPSR. F, forward; R, reverse; C1, CRISPR1.
